# Optimizing the *In Vitro* Neuronal Microenvironment to Mitigate Phototoxicity in Live-cell Imaging

**DOI:** 10.1101/2025.08.04.668446

**Authors:** Cassandra R. Hoffmann, Simon Maksour, Jordan E. Clarke, Maciej Daniszewski, Fiona J. Houghton, Jingqi Wang, Alice Pébay, Paul A. Gleeson, Mirella Dottori, Ellie Cho, Andrew Zalesky, Maria A. Di Biase

**Author notes:** Corresponding author: Cassandra Hoffmann.

## Abstract

Long-term imaging formats are ideal for capturing dynamic neuronal network formation *in vitro,* yet fluorescent techniques are often constrained by the impact of phototoxicity on cell survival. Here we present a live-imaging protocol that was optimised via quantitative analysis of 3 target culturing conditions on neuromorphological health: extracellular matrix (human- versus murine-derived laminin), culture media (Neurobasal™ versus Brainphys™ Imaging media), and seeding density (1×10^5^ versus 2×10^5^ cells/cm^2^). A cortical neuron reporter line was differentiated from human embryonic stem cells by transduction of Neurogenin-2 and green fluorescent protein, then fluorescently imaged in 8 different microenvironments daily for 33 days. Alongside viability analysis by PrestoBlue assay and gene quantification by digital polymerase chain reaction, an automated image analysis pipeline was developed to characterise network morphology and organisation over time. Brainphys™ Imaging medium was observed to support neuron viability, outgrowth, and self-organisation to a greater extent than Neurobasal™ medium with either laminin type, while the combination of Neurobasal™ medium and human laminin reduced cell survival. Further, a higher seeding density fostered somata clustering, but did not significantly extend viability compared to low density. These findings suggest a synergistic relationship between species-specific laminin and culture media in phototoxic environments, which is positively mediated by light-protective compounds found in Brainphys™ Imaging medium.

## Introduction

Live-cell imaging offers a time-resolved window into neuron morphogenesis. Applications of longitudinal imaging formats – spanning days to weeks – have long served investigations of progenitor differentiation^1–5^, neurodevelopmental and neurodegenerative disease phenotypes^6–9^ and treatment response *in vitro*^10^. Extended observational windows are particularly well-poised to capture the process of neuron network maturation, whereby neurites fasciculate into bundles and somata aggregate into clusters to facilitate optimal global communication^11^. Fluorescence microscopy represents a useful tool in this context due to its unparalleled signal quality and target selectivity, and offers a wide range of probes that include genetically-encoded protein reporters and membrane-permeant dyes. However, with few notable exceptions^12^, the extensibility of whole-cell live fluorescent imaging has generally been limited to periods of no more than two weeks ^8,10,13–19;^ beyond which more localized fluorescent tagging of biomolecules or organelles ^7,20,21^, or completely label-free methods^2,11^ are favoured. This paucity can be attributed to known complications of phototoxicity and photobleaching, which impact cell physiology and compromise signal clarity in a cumulative manner^22^. Optimising the *in vitro* cell microenvironment to maintain structural, metabolic, and trophic support throughout live-cell paradigms has therefore become imperative to fully harness the insights they offer.

Phototoxicity causes ultrastructural damage to cells by disrupting mitochondrial function^23^, lysosomal membrane stability^24^, and other biomolecular pathways^25,26^. Mechanistic inquiries have discerned reactive oxygen species (ROS) as a key contributor, with free radicals being produced by both light-irradiated cells and media^23^. Although classic media systems such as Neurobasal™ Plus with B-27 (henceforth referred to as NB medium) contain some antioxidant enzymes^27^, other speciality photo-inert media products have been designed to actively curtail ROS production with a rich antioxidant profile and the omission of reactive components such as riboflavin. The Brainphys™ Imaging medium with SM1 system (henceforth referred to as BPI medium) is one such example shown to protect mitochondrial health of neurons following violet light irradiation and exogenous hydrogen peroxide exposure^28^, and has been utilised in live-imaging applications to maintain cell health and improve fluorescent signal^29,30^. Its noteworthy performance in supporting electrophysiological maturation and synaptogenesis over longer periods of 2-3 weeks^28^ renders BPI medium an ideal candidate for extended imaging in phototoxic environments.

Alongside culture media, cell density has also emerged as a plausible modulatory target for mitigating the effects of redox imbalance. Sparse cultures are more vulnerable to pro-apoptotic mediators in general^31,32^, but show particular sensitivity to free radicals^33–35^. Conversely, high-density configurations confer shortened intercellular distances that are optimal for cell-to-cell exchange of protective neurotrophins, cytokines, and peptides ^36,37^. Corroborating evidence shows that high-density neuron cultures can survive without extrinsic supplementation of neurotrophins, whereas low-density populations lack the autocrine and paracrine functions to self-sustain under these conditions^38,39^.

A third consideration is the maintenance of extracellular matrix (ECM) coatings, which provide anchorage and bioactive cues for cell migration, behaviour, and differentiation^40^. In particular, the combination of synthetic ECM polymers, such as Poly-D-Lysine (PDL), together with purified biological ECM proteins, such as laminin, have been shown to synergistically promote neuron adherence while still allowing motile self-organisation^41–43^. Members of the laminin family exhibit further specialization based on their heterotrimeric α, β and γ chain constitution. While isoforms such as LN111, LN521, LN332, and LN511 maintain stem cell pluripotency and self-renewal^44,45^, LN511 seems to play an additional role in driving morphological and functional maturation of differentiated neurons^46–48^. LN511 is one of the most ubiquitously expressed isoforms throughout nervous system development^49^, with *in vivo* gene knockout studies implicating constitutive chains α5, β1, and γ1 in healthy neural tube formation^50^. Cell culture applications broadly utilise laminin isoforms from murine origin^51–53^, however completely xeno-free paradigms are now feasible with the commercialisation of human-derived laminins^48,54^. Comparative studies have demonstrated equivalent^55^ or superior^48,54^ functional development of neurons plated on human-derived laminin to its mouse-derived counterpart. Thus, relatively elementary adjustments to culturing protocols, such as optimizing extracellular matrix, increasing cell seeding density and improving media composition, have the potential to protect neuron viability and physiological maturation under phototoxic conditions.

Previous explorations of laminin, seeding density, or medium type on neuron health have largely centred on endpoint assays and immunostaining methods rather than lifespan microscopy. Here, we aimed to determine the synergistic effects of medium (NB or BPI), laminin (murine- or human-derived), and cell density (1×10^5^ or 2×10^5^ per cm^2^) in prolonging cell viability and promoting structural network organisation under longitudinal fluorescence imaging. To achieve this, we differentiated human embryonic stem cells into neurons by transducing with Neurogenin-2 (NGN2) and green fluorescent protein (GFP), then compared survival and morphogenesis in eight unique microenvironments over 33 days of once-daily scanning. To characterise cell morphology, we developed an automated and scalable bioimage analysis pipeline that quantified structural indicators of health, maturation, and complexity.

## Methods

### Human Embryonic Stem Cell (hESC) Culture

This study utilized a H9 hESC line (WA09, WiCell, hPSCreg ID: WAe009-A, normal XX karyotype^56^) that was previously shown to express pluripotency markers by Maksour et al^57^. Cells were maintained in TeSR-E8 (StemCell Technologies, #05990) or mTeSR1 (StemCell Technologies, #85850) on vitronectin XF (10 µg/mL, StemCell Technologies, #7180)-coated T25 flasks (#156800, ThermoFisher Scientific), at 37 °C with 5% CO_2_. At 60-70% confluence, cells were passaged using 0.5 mM EDTA (Life Technologies, #AM9260G) in phosphate-buffered saline (PBS; #14190250, Life Technologies) at a 1:20 ratio. Cells were passaged a total of 89 times. Media was changed every day.

### Lentiviral Production

In order to differentiate neurons from stem cells, viral particles with an open reading frame of NGN2 were generated using a protocol by Maksour et al^58^. Briefly, HEK293T cells were maintained in DMEM/F12 with Glutamax (Gibco, #10565-018) and 5% Foetal Bovine Serum (FBS; SFBS-F, Interpath) and passaged using Accutase (00-4555-56, Life Technologies). Twenty-four hours before transfection, cells were dissociated with Accutase for 2 mins and 4 x 10^6^ cells were seeded per T75 flask. Cells were transfected with the packaging plasmids vSVG (Addgene, USA, #8454), RSV (Addgene, #12253), pMDL (Addgene, #12251), and either the doxycycline-inducible lentiviral vector pLV-TetO-hNGN2-eGFP-Puro (#79823, Addgene) or the reverse tetracycline transactivator vector FUW-M2rtTA (#20342, Addgene). This was achieved by incubating plasmid DNA with Polyethyleneimine (Sigma-Aldrich, USA, #408727) at a ratio of 1:3 for 6 mins, then adding to cells in a ratio of 4:2:1:1, transfer vector:pMDL:RSV:vSVG. The viral supernatant was collected at 24, 48 and 72 hours, and viral particles were filtered with a 0.45 µm syringe filter. Virus was concentrated by ultracentrifugation at 66,000 × *g* for 2 hours at 4°C, resuspended in PBS, and cryopreserved at -80°C.

### Neuronal differentiation

Several methods can be utilized to direct stem cells to a neuronal fate. Extrinsic factors such as small molecules and growth factors may be used to recapitulate stages of neurogenesis^59,60^, while transcription factor overexpression^61,62^ may be used to program neuronal lineage, and other molecules^63,64^ to upregulate pro-neuronal genes. In the present study, cortical neurons were differentiated using a modified protocol by Maksour et al^58^ that combines NGN2 overexpression and developmental patterning. Single hESCs were seeded at 10,000 cells/cm^2^ on plates coated with Poly-D-Lysine (PDL, 10 µg/mL, P6407-5MG, Sigma-Aldrich) and mouse laminin (10 µg/mL, #23017015, Gibco, ThermoFisher Scientific) in TeSR-E8 medium supplemented with 10 µM Y27632 (#72304, Stemcell). Cells were infected with 0.5-1 µL of both pLV-TetO-hNGN2-eGFP-PURO and FUW-M2rtTA for 17 hours. After removal of virus particles, cells (denoted Day 1 cells) were differentiated in induction medium (Neurobasal™ Plus medium [#A3582901, Gibco, ThermoFisher Scientific] supplemented with GlutaMAX [100X, 10 µl/mL, #35050061, Gibco, ThermoFisher Scientific], B-27 [50X, 20 µl/mL, #17504044, Gibco, ThermoFisher Scientific], N2 Supplement [100X, 10 µl/mL, #17502048, Gibco, ThermoFisher Scientific], Insulin-Transferrin-Selenium-Sodium Pyruvate [ITSA, 100X, 10 µl/mL, #51300044, Gibco, ThermoFisher Scientific], LDN-193189 [10 µM, #04-0074, Stemgent], doxycycline hyclate [1 µg/mL, DOX; #D9891, Sigma-Aldrich], brain-derived neurotrophic factor [BDNF, 10 ng/mL, #78005, StemCell Technologies], and Y27632 [10 µM]). Day 3 cells were purified in selection medium (induction medium with 5 µM Y27632 and 2 µg/mL Puromycin [#73342, StemCell Technologies]). Cells were differentiated until Day 4, at which point they started to exhibit morphology consistent with neural progenitor identity, and were harvested by incubating with TrypLE (#12605010, Gibco, ThermoFisher Scientific) for 3 mins at 37°C. These were cryopreserved at -196°C in freeze down media (10% DMSO [#D2650, Sigma-Aldrich], 40% knockout serum replacement [#10828028, ThermoFisher Scientific], 50% neural media supplemented with 10 ƒM Y27632).

### Experimental conditions

Day 4 neurons were cultured in experimental conditions on 7 x 8-well ibidi untreated glass-bottom high chamber µ-Slides (#80807, DKSH) across 3 separate experiments. Three additional control chamber slides were maintained in an incubator with no live-cell imaging. All chamber slides contained one experimental condition per well (plate map presented in Supplementary Figure 1). These 8 conditions systematically tested all combinations of 3 factors: medium (NB vs. BPI), laminin (mouse vs. human) and seeding density (2×10^5^ vs. 1×10^5^). Cells were mycoplasma tested with the MycoAlert Mycoplasma Detection Kit (Lonza, #LT07-318).

To plate cells for conditions, chamber slides were coated with 10 µg/mL PDL in PBS and incubated at room temperature (RT) for 30 mins, washed twice with PBS and incubated with 10 µg/mL target laminin (mouse [#23017015, Gibco] or human [#LN511-0502, Biolamina]) overnight at 5°C. Laminin was then rinsed once with PBS, and 300 µl NB maintenance medium (Neurobasal™ Plus medium supplemented with GlutaMAX [10 µl/mL], B-27 [20 µl/mL], N2 [10 µl/mL], ITSA [10 µl/mL], DOX [1 µg/mL] and BDNF [10 ng/mL]) with 5 µM Y27632 was added to each well. Day 4 neurons were thawed in a water bath, collected slowly with 5 mL Neurobasal™ medium, centrifuged at 300 x g for 3 mins, and resuspended in 1 mL NB maintenance medium. A haemocytometer was used to seed cells at two target densities in monolayer format: 1 × 10^5^ cells per cm^2^ (denoted low density) or 2 × 10^5^ cells per cm^2^ (denoted high density). Although live-cell imaging studies generally employ sparser seeding^14,15^, these target densities were chosen based on pilot data showing rapid cell death at lower densities such as 0.5 × 10^5^ cells per cm^2^ (Supplementary Figure 2). Given previous literature has used high densities to enhance neuron survival in non-live cell imaging designs^65,66^, we sought to apply this strategy to a phototoxic context. Live-cell imaging commenced on Day 7 neurons (see next section). All neurons were maintained in NB maintenance medium supplemented with 1 µg/mL of target laminin until Day 14, whereby half of wells were switched to BPI maintenance medium (BrainPhys™ Imaging medium [#05796, STEMCELL Technologies] supplemented with NeuroCult SM1 [20 µl/mL, #05711, STEMCELL Technologies], N2-A [10 µl/mL, #07152, STEMCELL Technologies], GlutaMAX [10 µl/mL], ITSA [10 µl/mL], BDNF [10 ng/mL], ascorbic acid [200 nM, #72132, STEMCELL Technologies], and target laminin [1 μg/mL]). As the osmolarity and nutrient profile of BPI were designed for short-term imaging experimentation ^28^, BPI medium was introduced in 50% increments and gradually switched back to NB maintenance medium at Day 28 in line with manufacturer recommendations ^28^. All cells were maintained at 37°C and 5% CO_2_ and half medium changes were performed every 2-3 days.

### Live-cell imaging

Following the established utility of the Incucyte imaging system in monitoring neuron development longitudinally^21^, cells plated on 8-well ibidi µ-Slides were imaged from Day 7-40 with Incucyte SX5 and SX1 widefield platforms (Sartorius) inside an incubator supplied with 5% CO_2_ at 37°C. Sixty-four fluorescence images were collected per well once every 24 hours, each comprising one contiguous field of view (FOV) that together spanned the entirety of each well. Each image capture consisted of 300 ms acquisition time, totalling 19.2 seconds of non-overlapping captures per well. Images were taken with a 20X objective in the green channel (excitation: 441–481 nm, emission: 503–544 nm), with dimensions 1408 x 1040 pixels and pixel size of 0.62 μm. Incucyte images associated with this study are available in the public repository BioImage Archive at https://www.ebi.ac.uk/biostudies/bioimages/studies/S-BIAD1522. Control cultures were maintained in the same incubator, however, did not undergo live cell imaging.

### Immunofluorescent Staining and Confocal Imaging

Cultures were fixed in paraformaldehyde (#C004, ProSciTech) diluted to 40 µl/mL in PBS for 30 mins, then washed twice with PBS, permeabilized with 3 µl/mL Triton X-100 (#T8787, Sigma Aldrich) for 10 mins, and blocked with blocking buffer (100 µl/mL normal goat serum [#16210064, Life Technologies] in PBS) for 90 mins (all at RT). Cultures were then incubated with primary antibody prepared in blocking buffer overnight at 4°C, washed 3 times with PBS, and incubated with secondary antibody prepared in blocking buffer for 120 mins at RT. Primary antibodies utilized were 1:2000 rabbit anti-beta tubulin (#18207, Abcam) and 1:200 chicken polyclonal anti-MAP2 (#92434, Abcam). Isotype controls, used at matched concentrations, were rabbit IgG (#02-6102, Invitrogen, ThermoFisher Scientific) and chicken IgY (#50579, Abcam). Secondary antibodies were 1:500 goat anti-rabbit IgG conjugated with Alexa Fluor 488 (#A-11008, Invitrogen, ThermoFisher Scientific) and 1:350 goat anti-chicken IgY conjugated with Alexa Fluor 633 (#A-21103, Invitrogen, ThermoFisher Scientific). Nuclei were counterstained with Hoechst 33342 (#B2261, Sigma-Aldrich). Images were acquired with a Zeiss LSM900 confocal microscope with a 10x air objective (numerical aperture: 0.45). The pixel size was set to 0.62 µm, the scan time per pixel to 1.03 µs, and the optical section thickness to 4.2 µm. Alexa Fluor 488 was excited with a 488 nm diode laser and emitted between 505-645 nm, Alexa Fluor 633 was excited with a 640 nm diode laser and emitted between 645-700 nm, and Hoechst 33342 was excited with a 405 nm diode laser and emitted between 400-600 nm (all detected with a GaAsP detector). A z-stack was captured with 4.30 µm intervals, and the maximum intensity z-projection was computed. The same imaging parameters were used for experimental and isotype control cultures.

### CellROX Assay

Of all culturing factors tested in the live-cell imaging experiment, the interaction of human laminin with media type modulated cell viability to the greatest extent, and was thus targeted for reactive oxygen species analysis. Cells were plated at low seeding density and the assay was performed at the earlier timepoint of Day 30 to replicate human laminin conditions with negligible cell death. The CellROX Deep Red Reagent (#C10422, Invitrogen, ThermoFisher Scientific) was used to detect oxidative stress in cells via fluorescence microscopy. Analysed cultures comprised 3 live-imaged wells for each of the 2 conditions (BPI medium, human laminin and low seeding density; and NB medium, human laminin and low seeding density), 3 positive control wells (treated with the pro-oxidant compound Menadione^67^ at 100 µM for 30 minutes) and 3 negative control wells (no live-cell imaging). Live cells were incubated with 5 µM CellROX at 37°C for 30 minutes, then washed 3 times with PBS and fixed in PFA with Hoechst 33342 counterstain, as outlined in the *Immunofluorescent Staining and Confocal Imaging* section. Z-stack images were acquired with a Zeiss LSM900 confocal microscope with a 10x air objective (numerical aperture: 0.45). The pixel size was set to 1.25 µm, the scan time per pixel to 1.03 µs, and the optical section thickness to 4.2 µm. For all plates, CellROX was excited with a 640 nm diode laser at 2% intensity and emitted between 645-700 nm, and Hoechst 33342 was excited with a 405 nm diode laser at 0.7% intensity and emitted between 400-600 nm (both detected with a GaAsP detector). Four 2340 × 2626 × 46.2 μm FOVS were selected per well based on regions of highest cell density. For each FOV, CellROX signal was isolated within nuclei regions by calculating overlap with Hoechst 33342 mask. Sum intensity along the z-plane was calculated per pixel, then mean pixel intensity was calculated per image. To account for variability in cell density, CellROX intensity was normalized to nuclei count. Background correction was then applied by subtracting the normalized mean intensity of the negative control from the normalized mean intensity of each FOV. Nuclei counting was performed in Imaris (Version 10.2, Oxford Instruments) using the ‘Spots’ tool with XY diameter of 5.83 μm and z diameter of 12 μm, and all other processing was performed with MATLAB (version 2023a, Mathworks).

### PrestoBlue Viability assay

The PrestoBlue assay (#A1326, Invitrogen, ThermoFisher Scientific) was performed according to the manufacturer’s instructions on Day 42 cultures. Briefly, cells were cultured at 37 °C in 10% PrestoBlue viability reagent in NB maintenance medium for 40 mins. Supernatant was transferred to clear-bottom plates in triplicate. Fluorescence was measured with a FLUOstar Omega Fluorescence 556 Microplate reader (BMG LABTECH) at excitation 560 and emission 590 nm. Omega software and MATLAB was used to compare fluorescence readings between longitudinally imaged experimental cultures and non-imaged control cultures. All readings were corrected for background fluorescence by subtracting the no-cell control reading.

### RNA Extraction and Quantification

Day 42 cells were dissociated with TrypLE Express (#12604013, Gibco) and pelletized. RNA extraction was performed with the Qiagen RNeasy Mini Kit (#74104, Qiagen) according to manufacturer’s instructions. RNA was then quantified with the Qubit RNA Broad Range Assay Kit (#Q10210, Thermo Fisher) according to manufacturer’s instructions, and read by a Qubit 2.0 Fluorometer (Invitrogen). RNA concentration was out of range for two conditions that had undergone death events (human laminin, NB medium and low seeding density; and human laminin, NB medium and high seeding density) and so these were excluded from subsequent digital polymerase chain reaction (dPCR) analysis.

### dPCR

*SYN1* (encoding synapsin 1) and *DLG4* (encoding PSD-95) were selected for quantification based on their roles in functional maturation and connectivity^57,58^. The QIAcuity dPCR system (QIAGEN) was used with the QIAcuity OneStep Advanced Probe Kit (#250131, QIAGEN) in 24-well 8.5K nanoplates (#250011, QIAGEN). Each well contained 12 μl total reaction volume, comprising 3 μl OneStep Advanced Probe Master Mix, 0.12 μl OneStep Advanced RT Mix, 0.6 μl Taqman gene expression assay, 1.5 μl Enhancer GC, 0.35 ng template RNA (concentration based on previous optimisation) and RNAse free water. The selected FAM-labelled Taqman gene expression assays were Hs00199577_m1 (*SYN1*) and Hs01555373_m1 (*DLG4*). Sealed plates were processed in the QIAcuity instrument with a temperature profile of 1 x 40 min cycle at 50°C, 1 x 2 min cycle at 95°C, 40 x 5 sec cycle at 95°C followed by 30 sec cycles at 60°C and imaged in the green channel with an exposure and gain of 500 ms and 6 respectively. Absolute quantification was performed with the QIAcuity Software Suite 2.2.0.26, and each reading was normalized by GAPDH. Readings were taken from 3 plates that each corresponded to an independent experiment, with No Template Controls (NTCs) included for each gene. The SD of individual measurements resided below 30%.

### Bioimage analysis

Figure 1 presents an overview of morphological and network metrics used to quantify the impact of the 8 experimental conditions on cell structure. For each plate, each of the 64 Incucyte FOV images acquired per condition once daily for 33 days were processed separately: comprising a total of 118,272 images with 14,784 images per condition. The same processing workflow was used across all images. Image processing was completed with either the Spartan high performance computing system^68^ or computing systems equipped with an Intel Core i7-8550U processor (4 cores, 1.80 GHz) and 16 GB RAM, running Windows 10 Pro (64-bit) or an Intel Xeon Gold processor (16 cores, 3.6 GHz) and 256 GB RAM, running Windows 10 Edu (64-bit). Fiji^69^ software was used for preprocessing, which comprised converting raw 32-bit images into 16-bit, and rescaling image intensities to their min-max intensity range. The semi-supervised machine learning toolkit Ilastik^70^ was used to segment neurite and somata masks (Supplementary Figure 3), with all colour/intensity, edge, and texture pixel features selected for classifier training. Mean fluorescence intensity was determined by first identifying cell regions of interest by isolating areas of each pre-processed image that coincided with the Ilastik neuron segmentation mask. The mean pixel intensity of these modified images was then calculated. MATLAB and Fiji was used to calculate 5 structural metrics from segmented images. These included 3 morphological metrics to index cell growth (neurite length, branch points, and Disaggregation Index), and 2 network metrics to index self-organisation (Cluster Density Factor and neurite coherency). All metric data from longitudinal imaging sequences were normalized to the first day of imaging (Day 7). Of the morphological metrics, neurite length was calculated by skeletonising the neurite segmentation and summing the number of pixels. As skeletonization is a thinning operation that produces a single pixel-wide representation, pixel count directly confers neurite length. Normalised neurite branch points were calculated by summing the number of intersection points in the neurite skeleton, then normalizing to neurite length. The Disaggregation Index (DI) is a novel metric that was calculated to characterise the patterning of small somata structures with a balanced spread across the growth field (Supplementary Figure 4). It is described by,

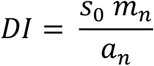

where s_0_ is the average somata area set by the user at the initial timepoint (here day 1 of imaging), m_n_ is the number of somata objects in the cell body mask (includes individualised and clustered cell bodies) at timepoint n, and a_n_ is the total area of all somata objects in the cell body mask at timepoint n. A high DI usually confers one of two cell configurations: young neurons that have not undergone clustering and remain individuated, or neurons that have clustered into groups that are relatively small and uniformly distributed. A low DI conveys a non-random cell configuration that usually signifies organised somata clustering, which occurs in comparatively mature cultures. In addition to morphological metrics, network metrics were developed following evidence that cytoskeletal networks with strong modularity – quantified by high somata clustering and neurite fasciculation^71^ – signify higher underlying functional maturity^72,73^. Here, the Cluster Density Factor (CDF) is a novel metric that was calculated to characterise the aggregation behaviour of somata (Supplementary Figure 4). It is described by,

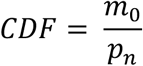

where m_0_ is the number of individual somata in the cell body mask at the initial timepoint, and p_n_ is the summed perimeter of somata objects in the cell body mask (includes individualised and clustered cell bodies) at timepoint n. A high CDF value indicates the presence of tight clusters with large numbers of somata condensed into smaller aggregates, which is a key hallmark of mature network formation. A low CDF value usually confers loosely clustered or spatially dissociated cultures.

The coherency metric was calculated to characterise the level of neurite fasciculation by quantifying angular alignment of neurites within a certain spatial proximity. The FIJI OrientationJ plugin developed by Fonck et al. in 2009^74^ was here implemented in a novel region-based configuration to examine neurites emanating from each somata or somata cluster. These sectional orientations were averaged for each somata object, which were in turn averaged across the entire image (workflow described in Supplementary Figure 5). To ensure computational tractability, the coherency metric was calculated for a subset of 9 representative FOV images per well, which were uniformly spaced across the entire well and kept consistent across all processing.

**Figure 1.**
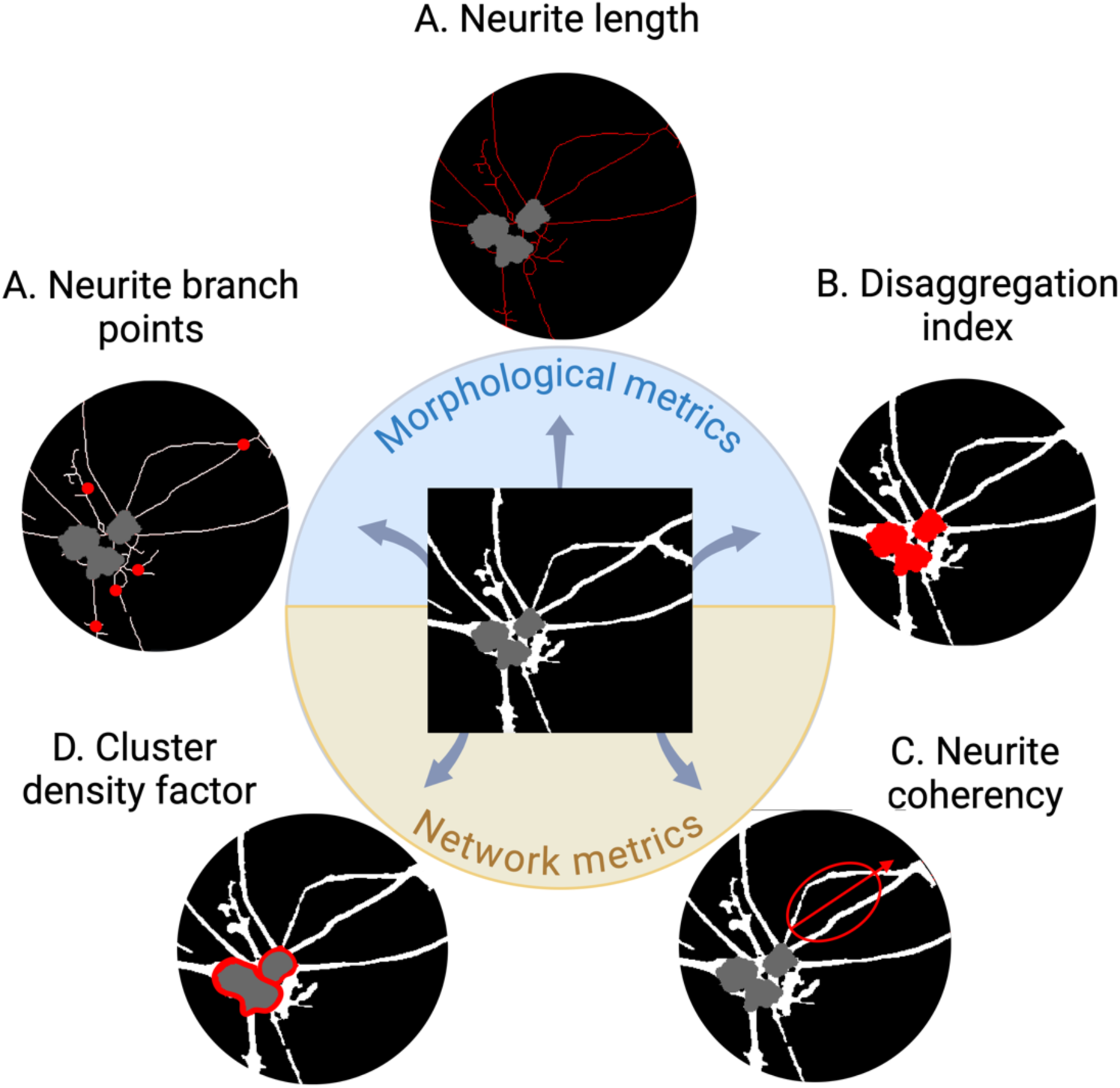
Morphological and network metrics. **(A)** Neurite length describes the number of pixels that comprise the one-pixel wide neurite skeleton. **(B)** Neurite branch points quantify the number of bifurcation and crossover points in the neurite skeleton. **(C)** The Disaggregation Index (DI) quantifies the uniformity of somata distribution across the growing field. **(D)** The Cluster Density Factor (CDF) describes the compactness of somata clusters relative to their perimetric size. **(E)** Neurite coherency describes the degree to which neurites are aligned in the same angular direction, which serves as an indicator of fasciculation.

### Statistical analysis

All statistical analyses were performed in MATLAB. For morphological analysis, cultures were considered non-viable and excluded from analysis at the timepoint when their average fluorescence intensity decreased more than 10% between successive measurements, denoted as the death event (example shown in Supplementary Figure 6). P-values were used to determine significance across all analyses, with significance level set to 0.05. Data was generated from 3 independent experiments (1 triplicate and 2 duplicates, totalling n = 7 per condition).

A timewise trajectory plot normalized by the initial reading was created for each of the 5 structural metrics (neurite length, branch points, DI, CDF, and neurite coherency) and fluorescence intensity. The AUC of the timewise trajectory was calculated to yield a time-averaged summary of each metric. Regression models were used to test the main effects and two-way interaction terms of the experimental conditions media (X_1_), laminin (X_2_), and seeding density (X_3_) on the AUC of each respective structural metric (Y), with plate effects included as a nuisance covariate. The interaction term model was specified as:

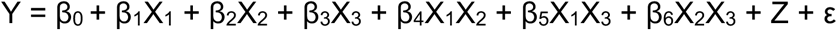

Where β_0_ denotes the intercept, β_1_, β_2_, β_3_, β_4_, β_5_, β_6_ are the coefficients for media, laminin, seeding density, media x laminin, media x seeding density, and laminin x seeding density respectively, Z represents the plate effect, and ε represents the error term. The Benjamini– Hochberg correction was used to correct for multiple comparisons, with all main effect and interaction term p-values included in the correction.

Network metric analysis was based on CDF and neurite coherency, since these two metrics capture somata and neurite behaviour throughout network formation. For each condition, both metrics were ranked according to their mean AUC, their ranks were added to form a summed rank, and summed ranks were used to calculate an overall network score per condition.

For CellROX ROS analysis, a linear mixed effects model was used to compare the effect of condition on normalized CellROX fluorescence intensity. Condition was modelled as a fixed effect while plate was modelled as a random effect, and FOV technical replicates were retained as observations nested within plates. The Benjamini-Hochberg correction was applied over 3 pairwise group comparisons (BPI vs. NB medium, BPI medium vs. positive control, NB medium vs. positive control). For PrestoBlue viability assay analysis, a Student’s t-test was used to compare experimental and control well means. Regression models with a within-subject factor for each well were used to test the main effects of each condition in control and experimental cultures.

## Results

### NGN2 Overexpression Directs hESCs to a Neuronal Fate

Upon completion of live cell imaging at Day 40, immunostaining revealed all cultures expressed neuronal marker class III beta-tubulin and dendritic marker microtubule-associated protein 2, indicating a differentiated neuronal phenotype (Supplementary Figure 7a-f). Phototoxic conditions had comprised cytoskeletal integrity to varying degrees across conditions, with the least blebbing and detachment evident in conditions with BPI media and human laminin (Supplementary Figure 7e,f). Furthermore, dPCR absolute quantification confirmed the expression of *SYN1* and *DLG4* in all analysed cultures, indicating the presence of both pre-synaptic and post-synaptic markers (Supplementary Figure 8).

### Neuron Viability is Compromised by Longitudinal Fluorescent Imaging

The PrestoBlue viability assay demonstrated that experimental cultures, live imaged daily for 300ms over the span of 33 days, exhibited lower viability than non-imaged controls in most conditions (Figure 2a,b). This indicates that phototoxicity likely played a role in cell death. Significantly lower viability was observed for experimental relative to control cultures plated on mouse laminin with BPI media and low seeding (t = -3.48, p = 0.0252); mouse laminin with NB media and high seeding (t =-15.25, p < 0.0001); mouse laminin with BPI media and high seeding (t = -3.71, *p* = 0.0206); human laminin with NB media and low seeding (t = -4.26, *p* = 0.0131); human laminin with NB media and high seeding (t = -3.27, *p* = 0.0305); and human laminin with BPI media and high seeding (t = -3.63, *p* = 0.0219). No significant difference was found between control and experimental cultures in mouse laminin with NB media and low seeding (t = -0.12, *p* = 0.9099), and human laminin with BPI media and low seeding (t = -0.39, *p* = 0.7126). Within control cultures, viability was increased in BPI media conditions relative to NB (t = 4.83, *p <* 0.0001); and high seeding density conditions relative to low (t = 2.62*, p* = 0.0108; Supplementary Figure 9a). No significant difference was found in laminin (t = -0.89, *p* = 0.3775). Within experimental cultures, viability was higher in mouse laminin conditions relative to human laminin (t = -3.84, *p* = 0.0003); and BPI media relative to NB media (t = 2.89, *p* = 0.0052; Supplementary Figure 9b). No significant difference was observed between seeding densities (t = 0.69, *p* = 0.4929).

**Figure 2.**
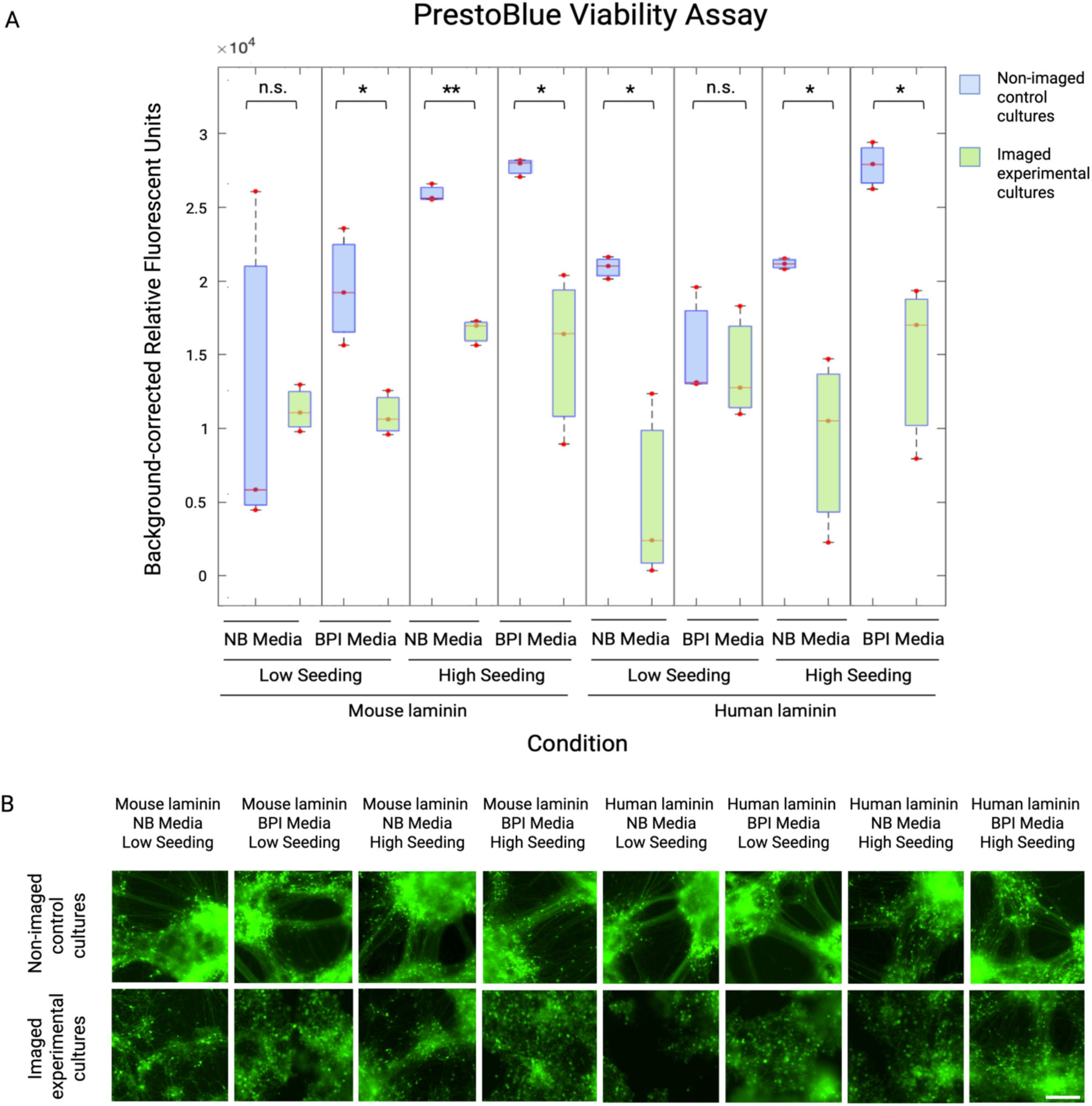
Viability analysis of Neurogenin-2 differentiated neuron cultures in non-imaged control and live-imaged experimental groups. **(A)** Student’s t-test analysis of PrestoBlue viability data revealed lower viability in experimental cultures (green boxes) compared to controls (blue boxes) in most conditions, specifically mouse laminin, BPI media, low seeding (1×10^5^ cells per cm^2^); mouse laminin, NB media, high seeding (2×10^5^ cells per cm^2^); mouse laminin, BPI media, high seeding; human laminin, NB media, low seeding; human laminin, NB media, high seeding; and human laminin, BPI media, high seeding. *p ≤ 0.05; **p < 0.0001; n.s. = no significant difference. Individual datapoints represent mean triplicates from the same well. Data generated from 3 experimental wells (n = 3 per condition) and 3 control wells (n = 3 per condition). **(B)** Control cultures that did not undergo any live-cell imaging (except a single endpoint scan at Day 42) maintained viability and cytoskeletal integrity to a greater extent than repeatedly imaged experimental cultures. Images acquired on Incucyte live-cell imaging system at Day 42. Scale bar = 200*μ*m.

### Culture Conditions Protect Neuron Architecture During Long-term Fluorescence Imaging

Timewise analysis of neurite length in live cell-imaged cultures revealed a characteristic biphasic trajectory of increased outgrowth peaking at Days 20-25, followed by a reduction in length for all conditions (Figure 3a). The declining outgrowth observed in the second phase could confer either apoptotic neurite retraction or healthy self-organisation^75^, and for this reason should be considered in the context of network-related metrics. The cultures grown in a combination of NB media and human laminin underwent early death events, whereas all remaining cultures self-organised into networks by Day 29 (Figure 3a,d). Neurite length area under curve (AUC) was significantly higher in the mouse laminin condition relative to human laminin (t = -5.67, *p* < 0.0001), BPI relative to NB medium (t = 6.71, *p* < 0.0001) and low relative to high seeding density (t = -4.44, *p* < 0.0001; Figure 3b). Isolation of the first developmental time period, from Day 7-18, revealed significantly higher neurite length in human laminin relative to mouse laminin conditions (t = 2.03, *p* = 0.0486) and low relative to high seeding density (t=-7.09, *p* < 0.0001), suggesting a more rapid onset of outgrowth in these conditions (Supplementary Figure 10a). Additionally, mean fluorescence intensity AUC was significantly higher in mouse relative to human laminin (t = -10.60, *p* < 0.0001; Figure 3c) across the entire cell lifespan. It was not significantly different for media types (t = 1.66, *p* = 0.1708) or seeding density (t = 1.15, *p* = 0.3389). Normalised neurite branch points were significantly higher in mouse relative to human laminin (t = -7.84, *p* < 0.0001), and BPI compared to NB media (t = 8.27, *p* < 0.0001) but not between seeding densities (t = -1.47, *p* = 0.2154), indicating higher neurite complexity in mouse laminin and BPI medium conditions (Figure 4a). Neurite coherency was significantly higher in mouse compared to human laminin (t = -7.04, *p* < 0.0001), BPI relative to NB media (t = 6.77, *p* < 0.0001), and high relative to low seeding (t = 2.78, *p* = 0.0190; Figure 4b). The DI, measuring the uniformity of somata across the growth field, was significantly higher in BPI relative to NB medium (t = 4.52, p < 0.0001). No significant difference was observed between seeding density (t = 1.98, *p* = 0.1144) or laminin (t = 1.44, *p* = 0.2149; Figure 4c). The CDF, measuring the level of somata clustering, was significantly higher in BPI media (t = 8.20, *p* < 0.0001) and high seeding density (t = 3.21, *p* = 0.0062), which confers a higher degree of somata aggregation in BPI and higher density conditions. CDF was not significantly different between laminins (t = -1.55, *p* = 0.1908; Figure 4d). Regression model *p*-values are summarised in Supplementary Table 1, timewise plots for morphological metrics in Supplementary Figure 10, and plate effects in Supplementary Figure 11.

**Figure 3.**
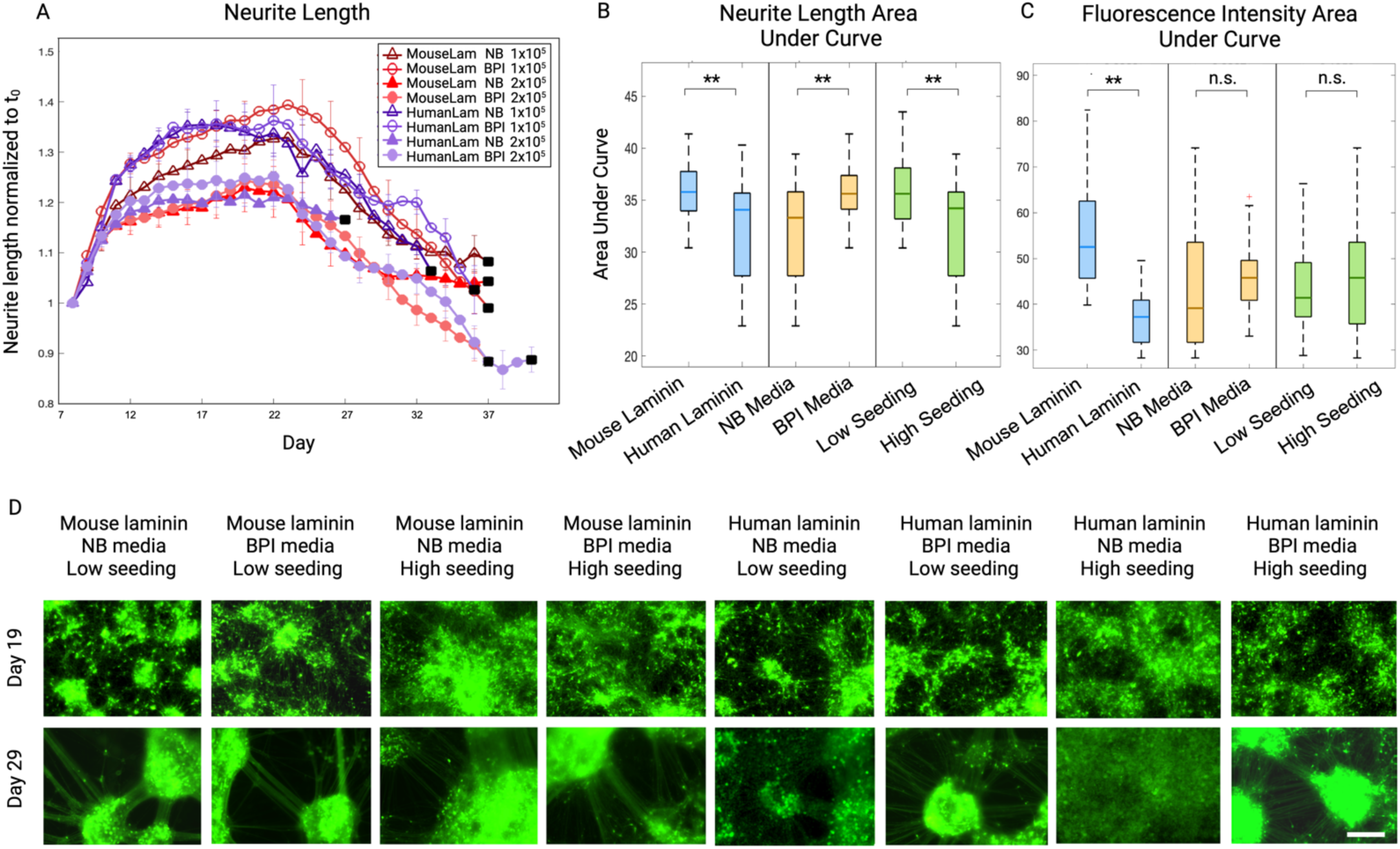
Morphological characterisation of Neurogenin-2 differentiated neurons live-imaged in 8 unique microenvironments. **(A)** A time series plot of neurite length in cultures imaged daily in the Incucyte system for 33 days, normalized to the initial timepoint (t_0_). Eight conditions compared the combinatorial effects of 3 factors: Neurobasal (NB) versus Brainphys Imaging (BPI) media, mouse versus human laminin, and low (1×10^5^ cells per cm^2^) versus high (2×10^5^ cells per cm^2^) seeding density. Plot datapoints represent mean of wells, and error bars represent standard error of the mean. Black squares demarcate cell death as defined by a >10% reduction in cell fluorescence between successive timepoints. **(B)** Boxplots present results of a regression model comparing neurite length across factors, based on cumulative area under curve (AUC) of time series plot. Cultures exhibited significantly increased neurite length in conditions featuring mouse laminin relative to human laminin, BPI medium relative to NB medium, and low seeding density relative to high density. Boxplot represents median and 25th and 75th quartiles, whiskers represent upper and lower values (*p ≤ 0.05; **p < 0.0001; n.s. = no significant difference). **(C)** Boxplots present results of regression model comparing mean fluorescence intensity AUC across factors. Cultures exhibited significantly increased mean fluorescence in conditions featuring mouse relative to human laminin, but no differences between NB and BPI media or low and high seeding conditions. **(D)** Incucyte live-cell images of neurons in each condition at Day 19 and 29. Cells in most conditions exhibited neurite outgrowth and self-organisation by day 29, except conditions featuring the combination of human laminin and NB media. Data generated from 3 experiments (n = 7 per condition). Scale bar = 200*μ*m.

**Figure 4.**
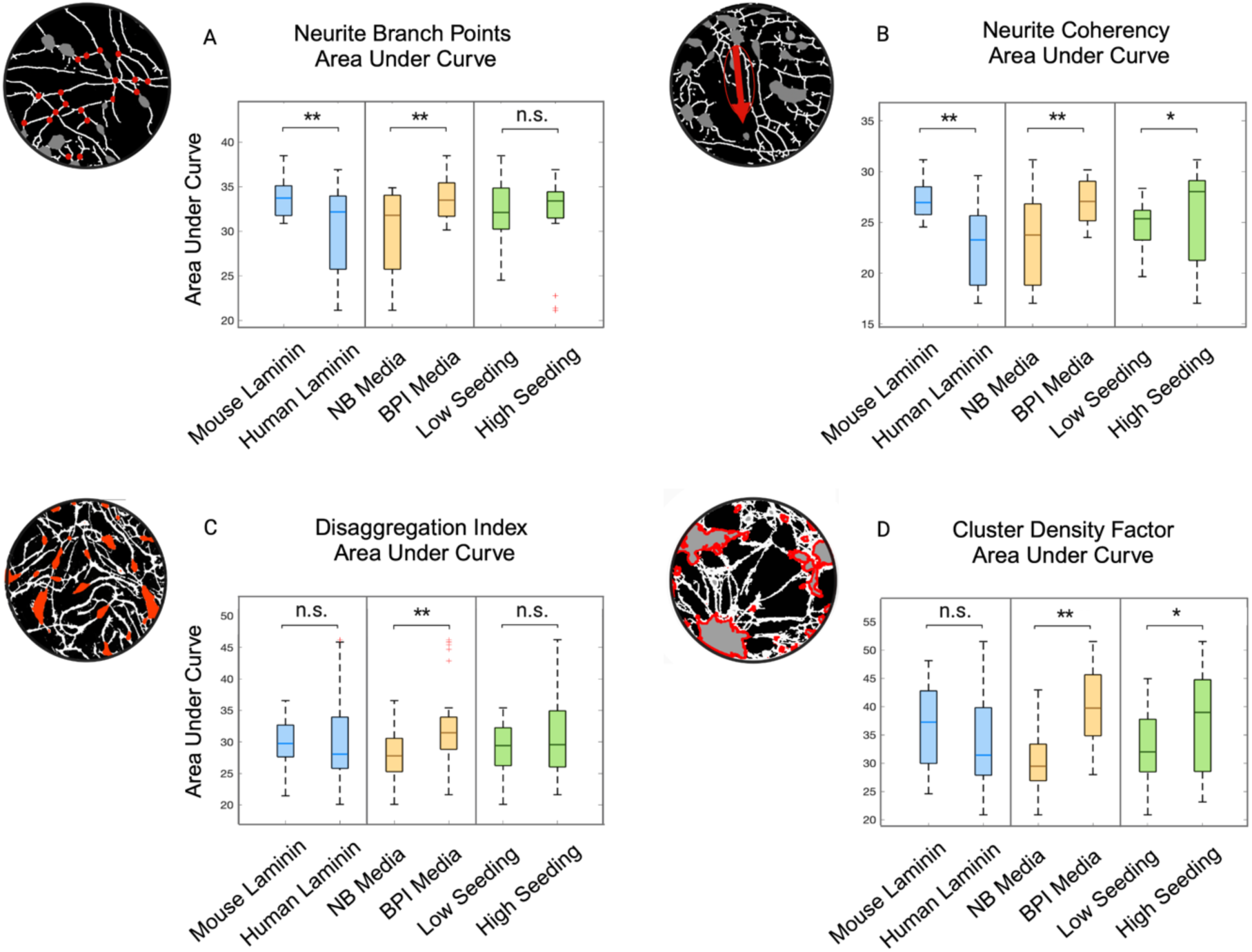
Structural quantification of Neurogenin-2 differentiated neurons cultured under different conditions comparing laminin, media, and seeding density. A bioimage analysis pipeline was established in Fiji to analyse morphological and network metrics of neurons. Boxplots for each metric present median and interquartile range of a regression model based on cumulative area under curve (AUC) of time series plot (*p ≤ 0.05; **p < 0.0001; n.s. = no significant difference, whiskers represent upper and lower values, red crosses represent outliers). **(A)** Neurite branch points were significantly higher in mouse relative to human laminin, and BPI compared to NB medium. **(B)** Neurite coherency was significantly higher in mouse compared to human laminin, BPI relative to NB medium, and high (2×10^5^ cells per cm^2^) relative to low (1×10^5^ cells per cm^2^) seeding density. **(C)** The Disaggregation Index was significantly higher in BPI relative to NB medium. **D)** The Cluster Density Factor was significantly higher in BPI relative to NB medium and high seeding density compared to low. Data generated from 3 separate experiments (n = 7 per condition).

In the regression model, a significant two-way effect was found between laminin and media on neurite length AUC (t = 5.10, *p* < 0.0001). While the impact of mouse laminin on neurite length was consistent across either media, human laminin combined with BPI medium, but not NB medium, had a positive effect on neurite length (Figure 5a,b). Accordingly, human laminin displayed significantly lower neurite length than mouse laminin within NB conditions (*p* < 0.0001), but not in BPI conditions (*p* = 0.7425; Figure 5a). No significant neurite length interaction effects were detected for laminin and seeding density (t = 1.86, *p* = 0.8025; Figure 5c), or media and seeding density (t = -0.46, *p* = 0.7745; Figure 5d). A significant two-way interaction effect was also found between laminin and media on fluorescence (t = 5.15, *p* < 0.0001), but not between laminin and seeding density (t = 1.85, *p* = 0.1217) or media and seeding density (t = -2.05, *p* = 0.1037).

**Figure 5.**
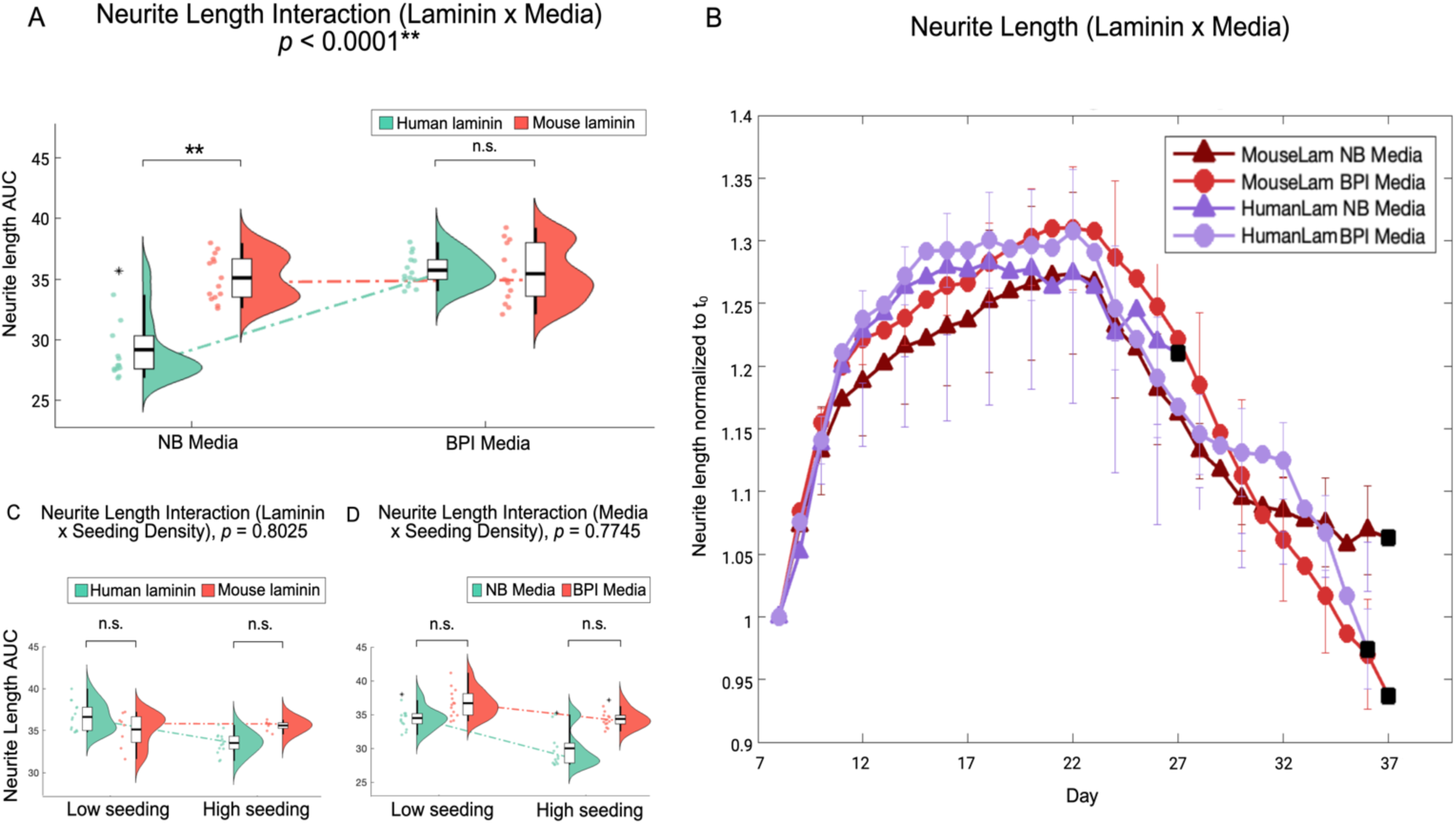
Media modulates the effect of laminin on neurite outgrowth in Neurogenin-2 differentiated neurons. **(A)** Results of a regression model reveal a significant two-way interaction effect between laminin type and media on the cumulative area under curve (AUC) of neurite length. Human laminin displayed significantly lower neurite length than mouse laminin within Neurobasal (NB) conditions, but not within Brainphys Imaging (BPI) conditions. (*p ≤ 0.05; **p < 0.0001; n.s. = no significant difference, violin plot represents mean, interquartile range, and data distribution as kernel density). **(B)** Timewise plots of laminin and media conditions across 33 days, normalised to initial timepoint (t_0_). The means of seeding density were calculated across different levels of media and laminin to isolate interaction effects. Error bars demarcate standard error of the mean, black squares demarcate cell death. **(C)** No significant interaction effects were detected for laminin and seeding density, **(D)** or media and seeding density. Data generated from separate 3 experiments (n = 7 per condition).

As presented in Figure 6, network metric analysis using CDF and neurite coherency revealed that both human and mouse laminin-plated cells cultured in BPI media at high seeding had the equally highest overall network scores, with the former displaying the overall greatest summed AUC. Conversely, human-laminin plated cells with NB media at both seeding densities ranked equally lowest for network formation (Supplementary Table 2).

**Figure 6.**
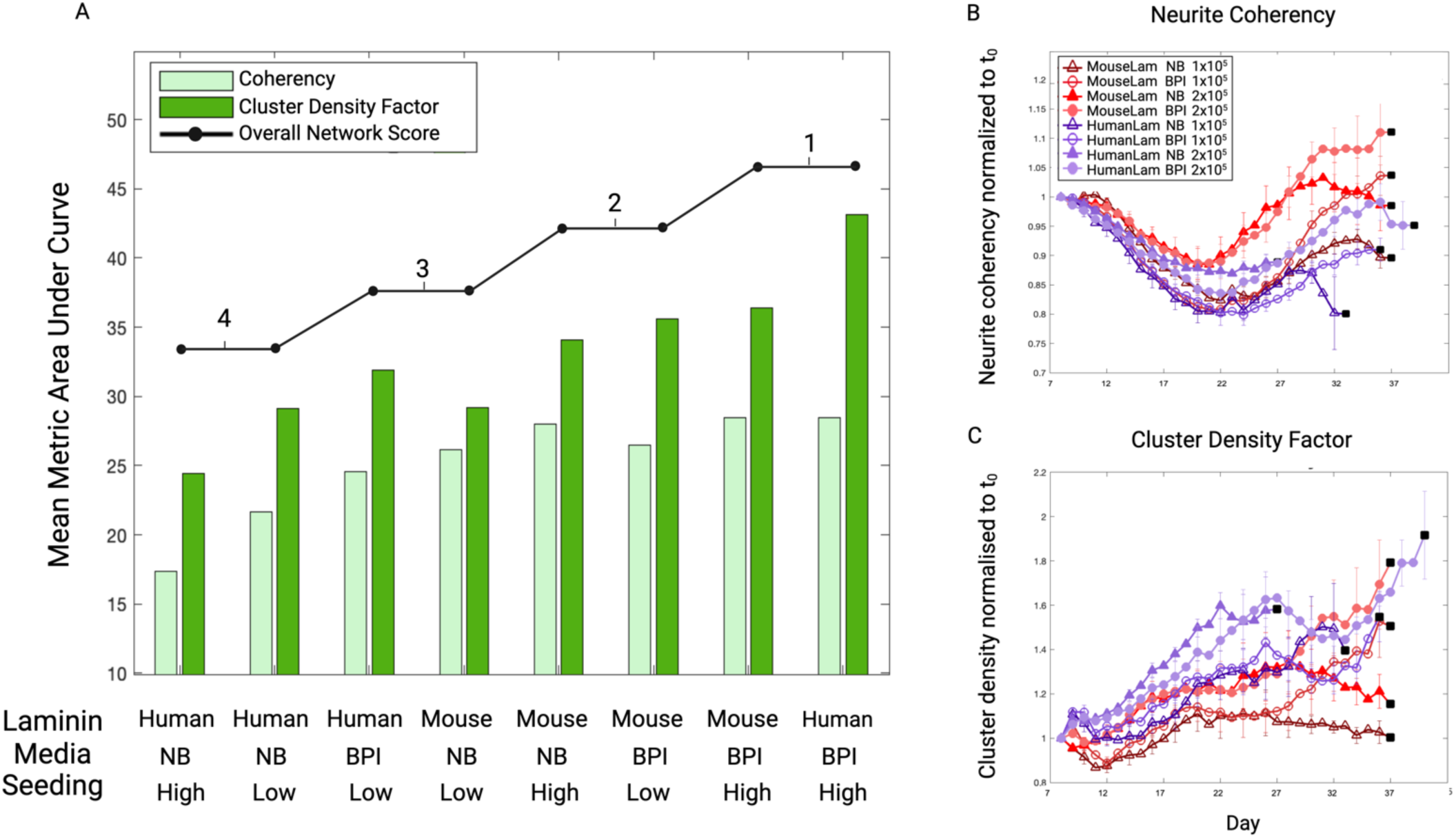
Culturing in Brainphys Imaging medium and at higher seeding density promotes more mature network formation. **(A**) Stem cell-derived neuron cultures were ranked based on somata clustering behaviour (conferred by Clustering Density Factor [CDF]) and neurite fasciculation (conferred by neurite coherency) area under curve (AUC) to give an overall network score. The two conditions with the tied highest score were human laminin with Brainphys Imaging (BPI) media and high (2×10^5^ cells per cm^2^) seeding density, and mouse laminin with BPI media and high seeding density. Human laminin with BPI media and high seeding density also displayed the highest summed mean AUC (Supplementary Table 2). **(B)** Timewise plot of coherency over the lifespan of the cultures (normalised to initial timepoint [t_0_]) reveals a trough around Day 22 where disorganised neurite outgrowth is at its greatest, followed by an upward trajectory. **(C)** Timewise plot of CDF over time reveals an overall upward trend as cultures organise. Plot datapoints represent mean of wells, and error bars represent standard error of the mean. Black squares demarcate cell death as defined by a >10% reduction in cell fluorescence between successive timepoints. Data generated from 3 separate experiments (n = 7 per condition). Low seeding density denotes 1×10^5^ cells per cm^2^.

### Reactive Oxygen Species are Central Mechanisms of Phototoxic Damage

Normalized CellROX fluorescence intensity was significantly lower in BPI conditions than NB (t = 2.7566, *p* = 0.0141; Figure 7), indicating reduced reactive oxygen species production. BPI medium-cultured cells also displayed significantly lower intensities than Menadione-treated positive controls (t = -3.2442, *p* = 0.0081), but there was no significant difference between NB-cultured cells and positive controls (t = -0.4875, *p* = 0.6291).

**Figure 7.**
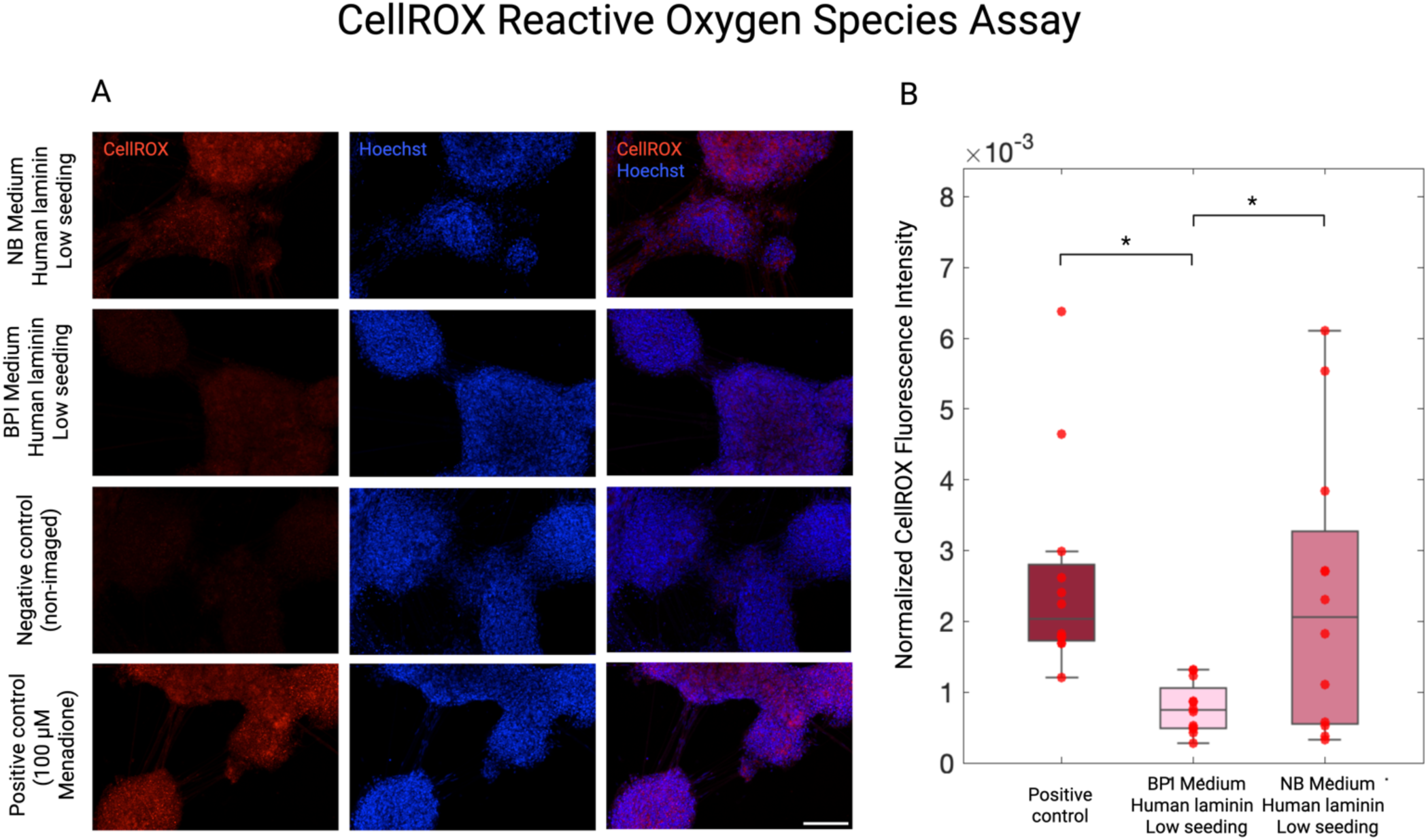
Reactive oxygen species underpin phototoxicity in neurons undergoing live-cell imaging. **(A)** The CellROX assay was performed on live-cell imaged neurons cultured in a Brainphys Imaging medium, human laminin, low seeding density condition; and Neurobasal medium, human laminin, low seeding density condition. Negative control cultures did not undergo live-cell imaging, and positive control cultures were treated with 100 µM Menadione for 30 mins. **(B)** Mean CellROX fluorescence intensity was normalized by nuclei count. Data was analysed with a linear mixed effects model with condition as a fixed effect and plate as a random effect to account for repeated within-plate technical replicates. *p ≤ 0.05. Datapoints represent FOV intensities. Data generated from 3 separate experiments (n = 3 per condition [n = 12 FOVs], n = 3 per positive control [n = 12 FOVs], n = 3 per negative control [n = 12 FOVs]). Scale bar = 150 *μ*m.

## Discussion

Although the longitudinal live-cell imaging format requires several methodological considerations, it is uniquely positioned to capture the entire lifetime of target morphogenic processes. One particularly valuable application of this is profiling neurodevelopmental and neurodegenerative disease progression. Indeed, live-cell imaging has already facilitated insights into the role of mitochondrial- and metabolic-associated genes in cellular respiration in autism models^76^, as well as early-stage cytoskeletal abnormalities in Alzheimer’s^9^ and Huntington’s^77^ disease models. The current study offers optimal culturing parameters to support and extend cell health in such applications, with a particular focus on culture medium, laminin type and seeding density. Morphological characterisation revealed that the use of Brainphys™ Imaging (BPI) medium in both mouse and human laminin-plated cultures supported neurite outgrowth, while the combination of Neurobasal™ (NB) medium and human laminin adversely impacted architecture. The most refined network organization was also observed in BPI media conditions, especially with human laminin at higher seeding density. By the end of experimentation, cell viability analyses revealed elevated survival rates in conditions incorporating BPI in contrast to NB medium, and mouse laminin in contrast to human laminin, but no difference across cell seeding densities. Based on these data, the following recommendations are proposed for future long-term live imaging protocols:

- BPI media should be used at high-volume fluorescent imaging periods to support neurite outgrowth and network formation. In line with manufacturer’s instructions, BPI media should be introduced incrementally after cells have spent minimum 7 days in differentiation medium (either Neurobasal™ or Brainphys™ systems) and should be used for a maximum period of 14 days^78^.
- In combination with BPI media, plates should be coated with either mouse or human laminin at a concentration of 10 µg/mL, and supplemented with 1 µg/mL laminin after 7 days in culture. Human laminin is optimal for studies of self-organisation and network formation, while mouse laminin should be favoured for longitudinal investigations requiring extended cell viability.
- Higher seeding densities such as 2 × 10^5^ cells per cm^2^ should be utilized in experiments that prioritize the development of modular neuron networks under live imaging conditions. However, sparser densities such as 1 × 10^5^ cells per cm^2^ may better serve morphological analyses requiring higher structural resolution for image analysis without loss of cell viability.

### Brainphys™ Imaging Medium and Neurobasal™ Medium

Culture medium is an attractive parameter for optimisation due to its regulatory role in cell health, identity, and morphology. BPI medium was formulated as an alternative to NB medium with the aims of mitigating light-induced neuronal damage and promoting synaptic maturation through optimal osmolarity, additives, and glucose concentration^78^. The results of the current study substantiated these original targets by revealing decreased ROS production and enhanced network morphology in light-irradiated cells cultured in BPI compared to NB media.

Historically, the Neurobasal™ B-27 media system has remained a common fixture in cell culture for over 30 years^27^ due to its established efficacy in maintaining long-term differentiated neuron culture^79,80^ while limiting glial proliferation^81^. The BPI system was subsequently formulated to support electrophysiological health in quantitative imaging applications, and has indeed demonstrated comparable performances to benchmark preparations (eg. artificial cerebrospinal fluid) in assessments of current-voltage profile, fast-rising calcium spiking, and network synchrony^28^. Given established multimodal evidence linking emergent neuron network morphology with electrophysiological maturity^71–73^ ^75,82,83^, our study sought to complement these functional data by tracking the structural self-organisation of cells. Our finding that cultures in BPI medium display greater outgrowth and network-like morphology justifies the use of this medium in future longitudinal live cell paradigms.

BPI’s protective capabilities arise from a lack of photo-reactive components and associated mitigation of oxidative stress. This has been established in previous literature^28^ and is recapitulated here as lower levels of ROS in BPI medium-cultured cells compared to NB medium. Conventional media additives such as riboflavin, while optimally supporting nutrient and pH requirements of cells, are potent photosensitizers that generate oxygen radicals when exposed to light. Upon absorption of a photon, these molecules transition from a ground state to a fluorescent singlet state, and can progress to a highly reactive triplet state that interacts with oxygen to produce ROS^75,76^. The omission of riboflavin in BPI media eliminates a significant pro-oxidative contributor, leaving a much smaller load from cellular metabolism^25^. Although it is recommended to reintroduce either NB or original Brainphys™ after a 14-day exposure period in order to replenish cells with photoreactive nutrients^28^, BPI media was released recently in 2020^28^ and emerging research may seek to build upon the manufacturer-recommended protocol.

### Mouse-derived and Human-derived Laminin

Laminin is routinely employed in 2D neuron culture because it provides a highly stable substrate for growth^84–86^, synaptogenesis^86^, and axon specification^87^, and offers a more defined composition than alternatives such as Matrigel^88^. While options such as human recombinant laminin are genetically engineered for optimal purity, others such as mouse laminin derived from Engelbreth-Holm-Swarm sarcoma retain high bioactive diversity due to their natural origin^89,90^. In the current study, we found a general trend of increased viability and neurite outgrowth in cells plated on mouse-derived relative to human-derived laminin; however human laminin conditions displayed a comparatively more rapid onset of neurite growth at the start of the imaging period. We noted that media type strongly modulated the health profile of cells within human laminin conditions, with cultures in BPI media performing at least equivalent, and in some cases superior, to those in corresponding mouse laminin conditions. Specifically, human laminin-plated BPI-cultured cells at high seeding density exhibited superior network formation in maturity, maintained cytoskeletal integrity until the end of experimentation, and ultimately outlasted all other conditions in lifespan. As BPI medium mitigates ROS production, these results may illuminate a distinct species-specific susceptibility of laminin to phototoxic conditions.

Laminins across animal species display differential properties based on their glycoprotein composition. The α subunit in particular has been shown to play a large role in determining these properties^91^, with human laminin α5 subtypes such as LN511 inducing different morphological and functional cell outcomes to murine-derived equivalents^48^. Integrins are the major laminin receptor class at the cell surface acting to drive this specification^91^. These receptors, being particularly concentrated at neuronal growth cones^92^ and synaptic junctions^93^, steer synaptic plasticity and stabilisation of the actin cytoskeleton^94^. Although not directly investigated here, evidence that laminin α5 harbours the most integrin binding sites of any subtype^95^, alongside established links between redox imbalance and the integrin-laminin interface^96^, favours the theory that integrin interactions may underlie the extensive death we observed in human laminin and non-ROS mitigating NB media combinations. Future research should seek to elucidate cell-ECM interactions mediating this process, and further, differentiate reactivity profiles of laminin origin species from laminin isoform. We chose to focus on murine-derived LN111 and human-derived LN511 based on their well-characterised suitability for neuron maintenance^46,58,97,98^, however, other investigations may benefit from comparing distinct isoforms from the same origin species. For example, human LN111 and LN511 exhibit different integrin binding behaviour – interacting with a narrower and broader range of sites, respectively^99^ – which likely manifest as divergent neuron adhesion and signalling outcomes.

### High and Low Seeding Densities

Cell-to-cell proximity is another factor that has been shown to modulate the effect of oxidative stress on culture^34^. Given the established neuroprotective effect of higher cell density^38,39^, our finding of increased neurite outgrowth in the lower seeding condition was unexpected. However, the accompanying decrease in cluster density relative to highly-seeded conditions suggests the observed reduction in neurite length reflects self-organisation rather than neurite retraction. Previous work has highlighted the tendency of dense cultures to undergo accelerated synaptogenesis and functional maturation, likely due to a higher proportion of successful connections resulting from axon pathfinding^100^. Corroborating work has established minimal plating densities necessary for neurons to generate healthy synchronised bursts and oscillations in culture^101^. In these formats, protective molecules such as neurotrophins and cytokines are secreted in maximal amounts ^36,37^, facilitating electrophysiological development and resilience to stress.

### Limitations and Considerations for Future Work

The work presented here demonstrated the significance of the cellular microenvironment in reducing cell death in phototoxic conditions; however, had a scope limited to 3 factors: media, origin of laminin, and seeding density. Other modifiable factors exist and should be considered in future research. Selecting proteins or dyes that fluoresce at longer light wavelengths in the red, far-red, or near-infrared range, such as mCherry, tdTomato, or infrared fluorescent proteins (iRFPs), generally confer less photodamage than their more energetic violet-shifted counterparts^102^. However, this advantage should be considered alongside the constraint of lower intensity and faster decay of fluorescence signal^103,104^, which is delimited primarily by a lower quantum yield of photons than GFP constructs^104,105^. Outside of the cellular microenvironment, microscopic imaging schemes fundamentally determine light irradiation parameters; tuning exposure time and excitation light power can mitigate cell phototoxicity by attenuating total dose or spreading lower doses over longer timeframes^106^. Of these, pulsed light schemes theoretically allow molecules occupying the singlet state to relax back into ground state rather than advancing to a ROS-inducing triplet state^107,108^. Supporting experimental evidence has been mixed, with some studies highlighting favourable outcomes on cell viability and physiology in pulsed compared to continuous laser illumination^109,110^, and others minimizing its impact in favour of direct antioxidant supplementation in media^111^. Entire integrated microscopy systems have also been developed to expressly minimise light exposure in live-cell imaging^112^. Light-sheet microscopy, especially lattice light-sheet, is one example that has the power to restrict extraneous light dose by illuminating only a single focal plane at a time^113^. Although direct comparisons of phototoxicity-mitigating techniques are lacking in the field, incorporation of as many phototoxicity-attenuating strategies as feasible, alongside assays to monitor the level of phototoxicity, are crucial in live-cell imaging design.

In the domain of quantification, the structural metrics developed here are well-suited for future examinations of monolayer neuronal culture, especially disease models created with patient-derived induced pluripotent stem cells^114^ or pharmacological agents^9,115^. Multidimensional characterisation of cell morphology assists in clarifying how genetic variants or environmental factors manifest as pathological phenotypes, which has clear implications for *in vitro* diagnostic and therapeutic research. However, the current pipeline has some limitations that could be further refined. First, our DI and CDF measurements require a baseline scan for comparison to subsequent timepoints, preventing the study of truly irradiation-naïve controls. Instead of using live cells, future studies could circumvent this limitation by analysing fixed and immunostained samples in treatment/control groups at each timepoint, employing a cross-sectional rather than longitudinal design. Second, our equations assume that cell count and size remain constant throughout the lifespan of the culture, specifically by the inclusion of s_0_ and m_0_ terms in DI and CDF calculations, respectively. This does not account for the fact that some degree of cell death is expected and inevitable under typical culturing conditions. As the primary focus of this work was to isolate the effects of phototoxicity under standardized fluorescent exposure, no secondary fluorescent viability dyes such as CytoTox^116^ were used to monitor live and dead cell proportions in real time. In future work, inclusion of these viability data could inform a dynamically updated term in DI and CDF equations along longitudinal image sequences, enabling more accurate timewise metric estimation. Third, the morphological network measures utilized here quantify structural neuron network morphology in isolation. Complementary analysis of functional network phenotypes – via techniques such as microelectrode array^117,118^ or calcium imaging^119,120^ – would lead to a more comprehensive understanding of neuron self-organisation. This is especially pertinent in examinations of BPI media, which has been shown to promote action potential generation and functional network synchrony^28^.

## Conclusions

Taken together, the results of this long-term imaging study demonstrate that neuron culture medium composition, extracellular matrix, and cell seeding density significantly impact cell viability and network formation in phototoxic environments. Specifically, the photo-inert formulation of BPI media extended cell viability and supported self-organisation by curtailing ROS production. A substrate of human laminin promoted network development when paired with BPI medium, while mouse laminin supported neuron longevity across medium systems. Finally, a lower seeding density increased neurite complexity, while a higher density fostered more sophisticated network modularity. Optimisation of these microenvironment features is pivotal in maintaining healthy neuronal structure and emergent network architecture during extended live-cell imaging experiments. Systematic characterisation of these parameters enables reliable longitudinal investigation of neuronal structure-function relationships in cell models of health and disease.

## Declarations

### Ethics approval and consent to participate

Informed consent was obtained for all embryo donors according to national and institutional guidelines. The study was approved by the University of Melbourne Research Ethics and Integrity Committee (LNR 4B) on 14/03/2024 as part of a project entitled ‘Schizophrenia hiPSC study 2’ (reference number 2024-28982-51093-4).

### Consent for publication

Not applicable.

### Availability of data and materials

The datasets generated during the current study are available in the BioImage Archive repository, https://www.ebi.ac.uk/biostudies/bioimages/studies/S-BIAD1522.

### Author’s contributions

Conceptualization, C.H., M.D.B., E.C., A.Z.; Investigation, C.H. and S.M.; Writing – Original Draft, C.H.; Writing – Review and Editing, C.H., S.M., J.E.C., M.D., F.H., J.W., A.P., P.A.G., M.D., M.A.B., E.C., A.Z.; Supervision, M.D.B., A.Z., E.C.; Visualization, C.H., A.Z., E.C.

## Acknowledgments

This research was supported by The University of Melbourne’s Research Computing Services, the Petascale Campus Initiative and Operational Infrastructure Support from the Victorian Government. In addition, the Victorian Centre for Functional Genomics provided access to the Incucyte SX5 system, with technical guidance from Dr Ada Koo. Furthermore, Vinod Dagar provided specialist advice for operation of the QIAGEN QIAcuity dPCR instrument.

The authors declare that they have not used AI-generated work in this manuscript.

