## Supplementary files for "Optimizing the *In Vitro* Neuronal Microenvironment to Mitigate Phototoxicity in Live-cell Imaging"

**Supplementary Material**

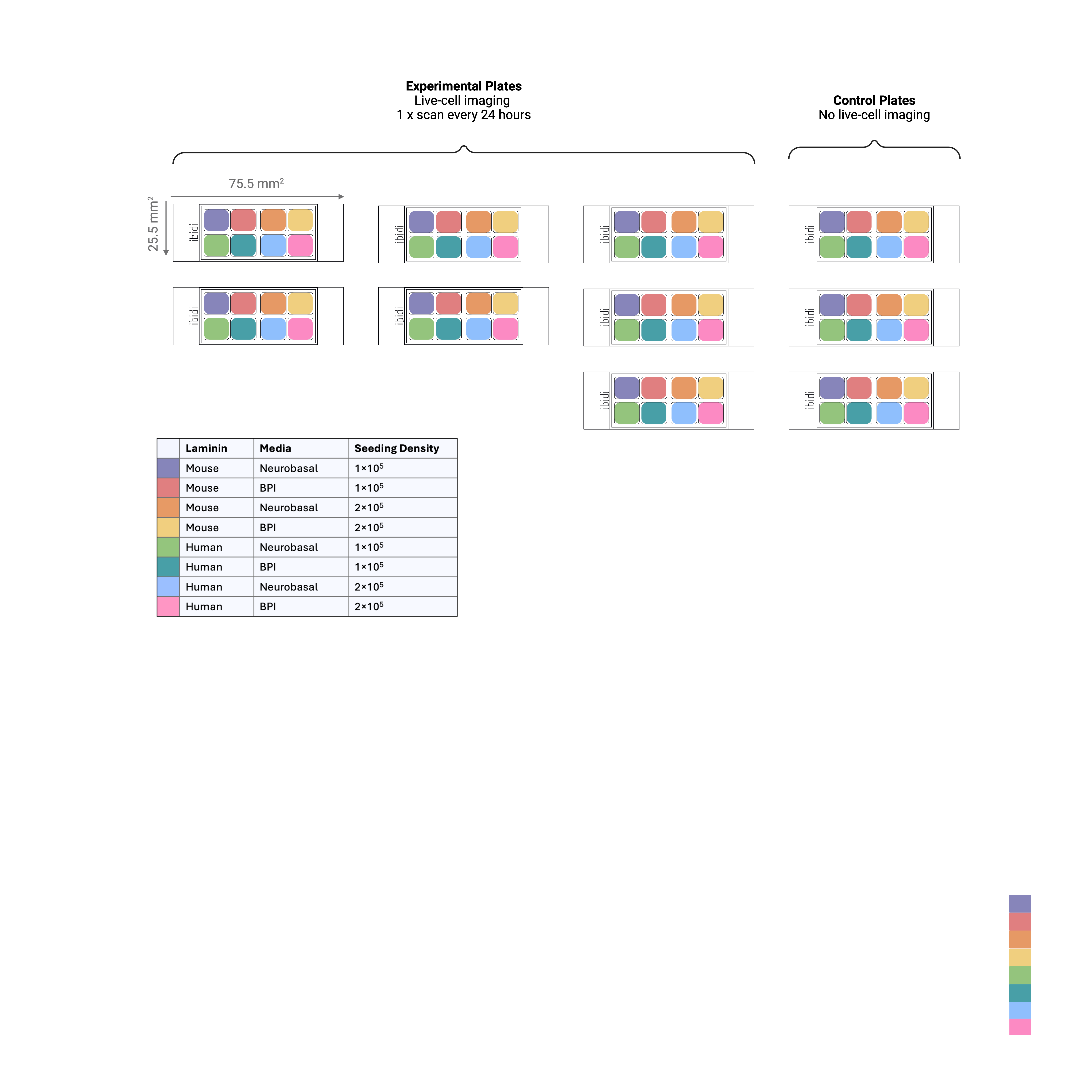

**Supplementary Figure 1. Experimental design.** Each plate contained 8 conditions that systematically tested all possible combinations of medium (Neurobasal^TM^ or Brainphys^TM^ Imaging [BPI]), laminin (murine- or human-derived), and cell density (2×10^5^ or 1×10^5^ per cm^2^). Three experiments with a total of seven plates were tested in the Incucyte live-cell imaging system. Three control plates were also maintained in a normal incubator with no ongoing imaging. Plate dimensions were 25.5mm^2^ x 75.5mm^2^.

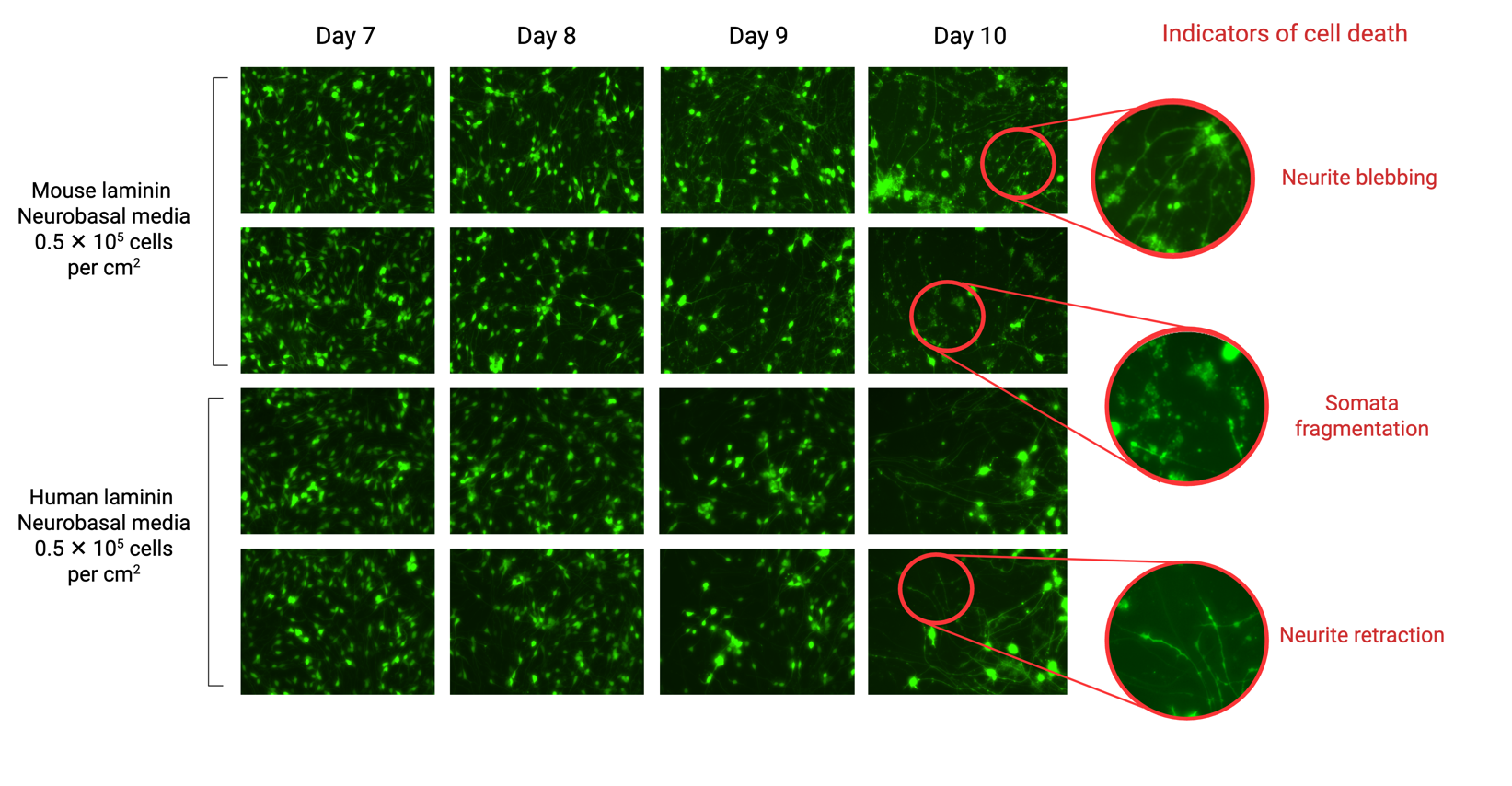

**Supplementary Figure 2. Live-cell imaging of lower seeding densities (0.5 × 10^5^ cells per cm^2^) induced rapid cell death, indicated by neurite blebbing, somata fragmentation, and neurite retraction.** This effect was observed across both laminin types in Neurobasal conditions, and occurred before the manufacturer-recommended timepoint for Brainphys Imaging medium introduction (Day 14).

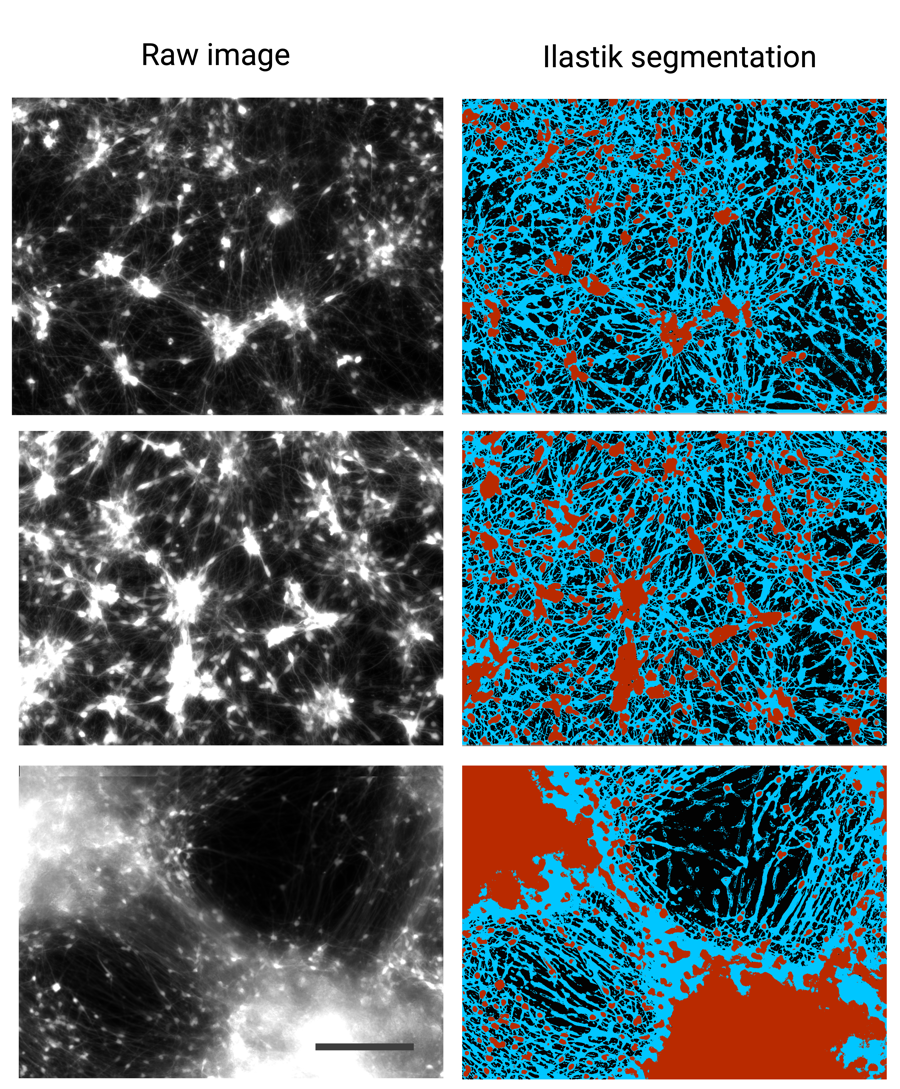

**Supplementary Figure 3. Raw images and corresponding Ilastik segmentations.** Somata masks are represented in red, and neurite masks are represented in blue. Scale bar = 200𝜇m.

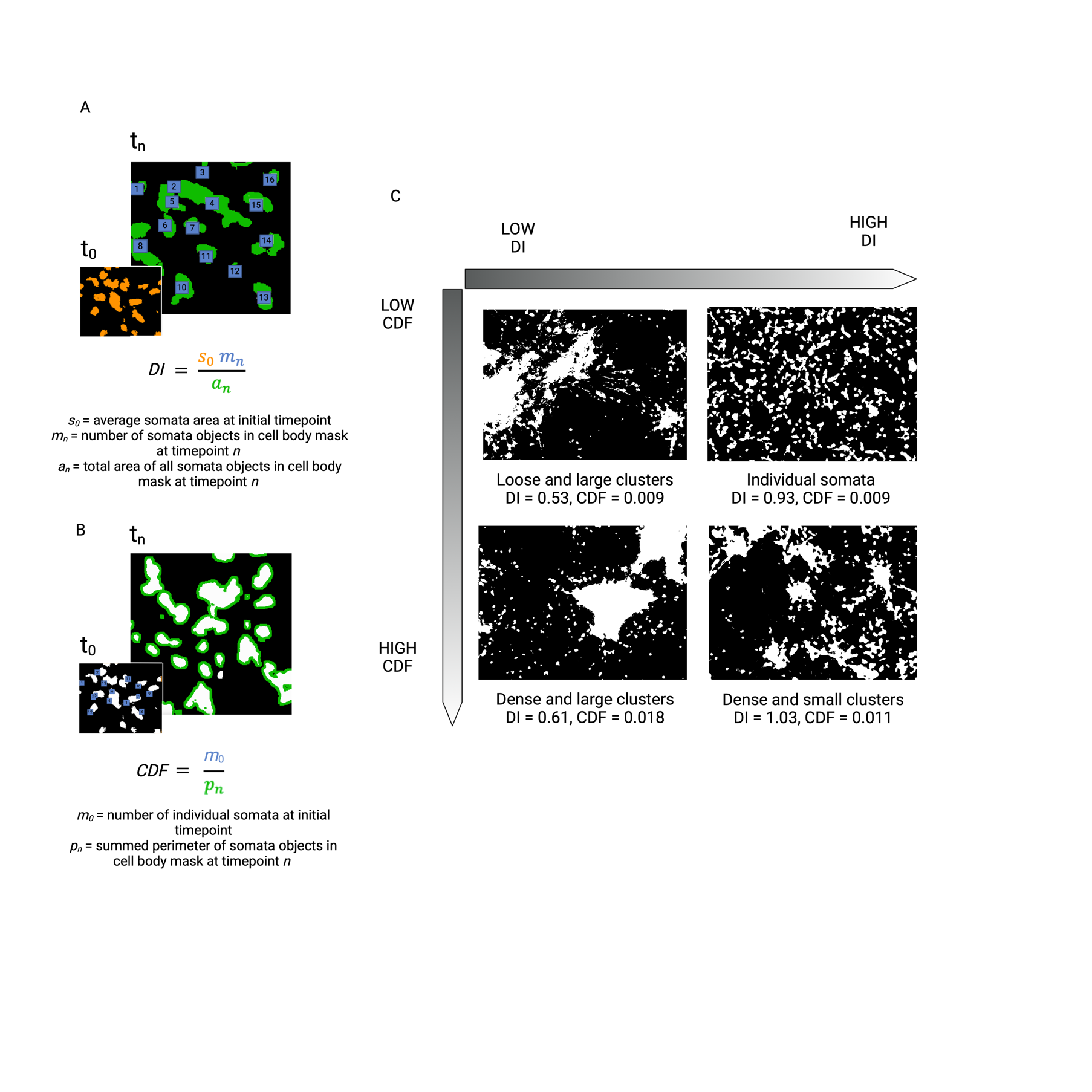

**Supplementary Figure 4. Calculation and examples of somata metrics. (A)** The Disaggregation index (DI) captures the degree to which somata are individuated or localized to small groups. The DI equation compares the total somata area at timepoint *n* to the initial timepoint, before organised cell migration has taken place. At the start of processing, the user determines the average somata area s_0_ by defining a circular ROI representative of a typically-sized somata. **(B)** The Cluster Density Factor (CDF) captures the compactness of aggregated somata structures. The CDF equation is a ratio of the amount of somata at the initial timepoint to the total perimeter of somata at timepoint *n*. This metric reflects the fact that exposed cell surface area is reduced as cells pack into dense clusters. **(C)** A high DI usually signifies that cells are either highly individuated, or assembled into small and relatively uniform clusters. A low DI usually confers that cells have migrated into localized clusters. A high CDF usually represents more networked cultures where most somata are associated with dense clusters. A low CDF usually denotes cultures at early or intermediate points of development that have not undergone extensive self-organisation.

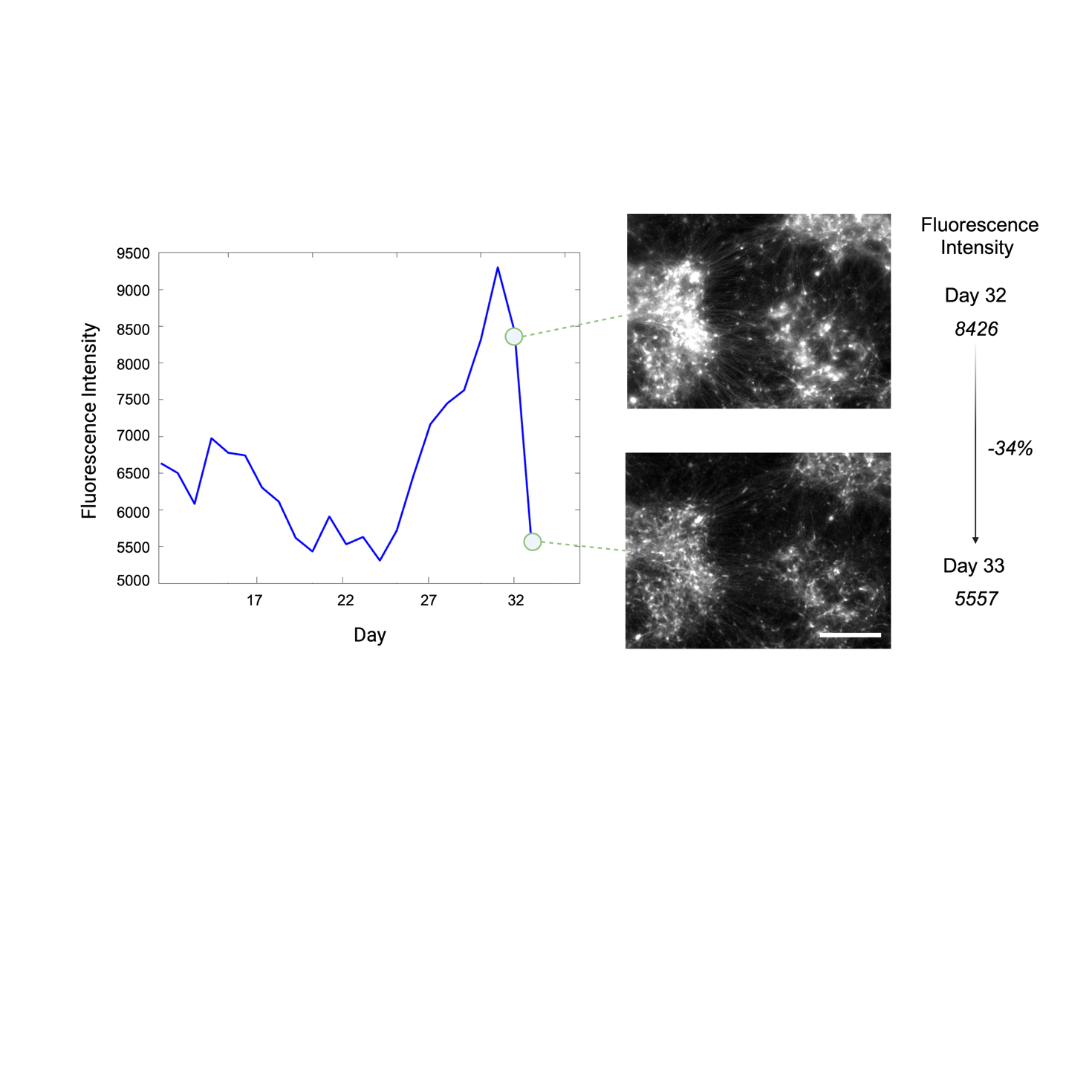

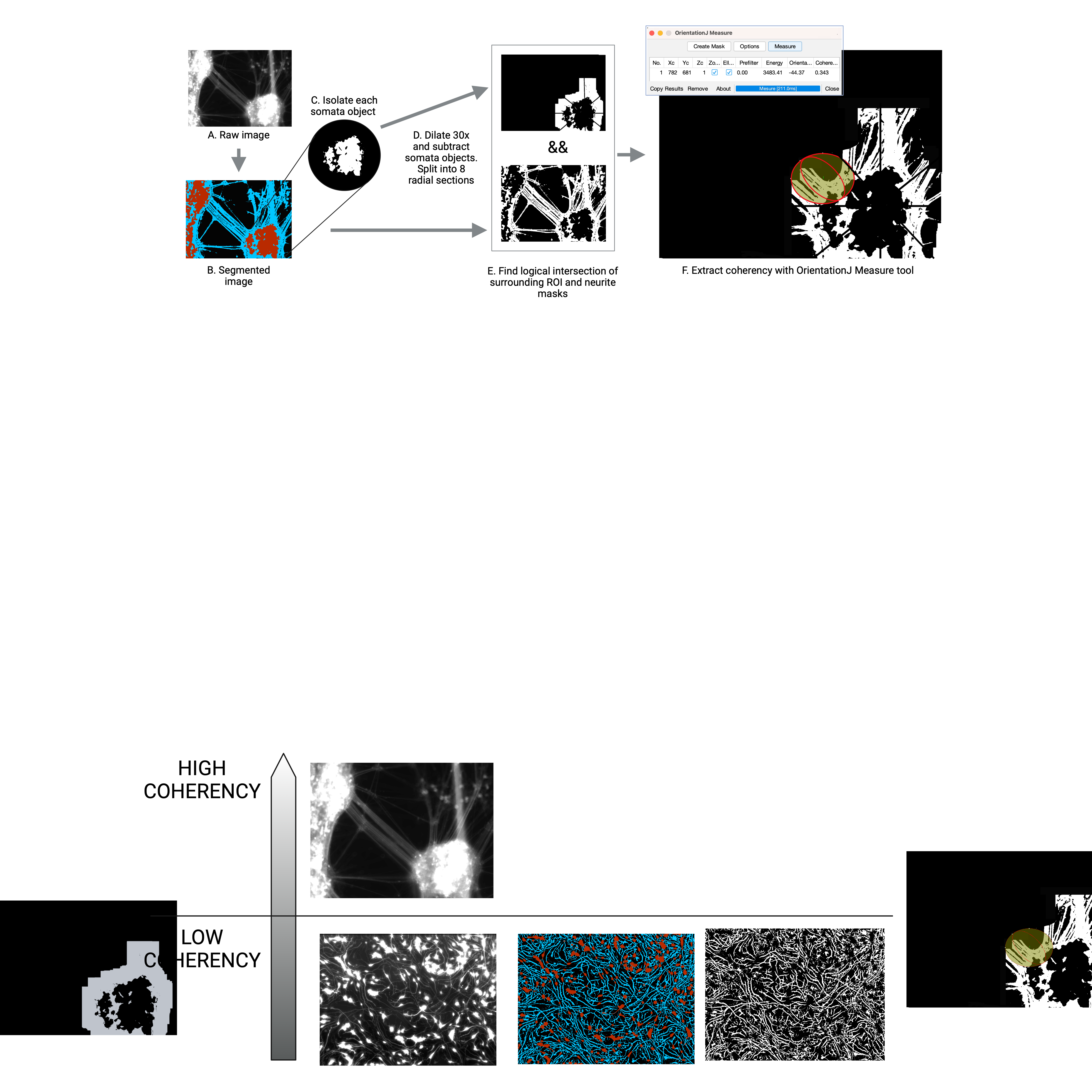
**Supplementary Figure 5. Workflow for calculation of neurite coherency. (A-B)** Raw images were segmented with Ilastik software. **(C)** Each object in the somata mask was isolated and processed separately to ensure intercellular neurite variability was captured. In this context, a somata object represented either an individual or cluster of cell bodies. **(D)** Each somata object in the cell body mask was dilated 30 times (~24.8μm) to create a surrounding region of interest (ROI). This dilation factor was selected to capture sufficiently sized neurite segments while minimising redundant, overlapping coherency readings with neighbouring somata structures. The surrounding ROI was sectioned into 8 radial segments for processing to minimise conflicting orientation measurements from orthogonal neurites. **(E)** Neurites within the ROI were isolated by finding the logical intersection of the surrounding ROI and neurite masks. **(F)** The Measure tool from the FIJI OrientationJ Plugin was used to measure the dominant orientation of neurites within each segment, with the width of the Gaussian kernel (sigma) set to 1.0. The mean coherency between 8 segments was calculated to create a score for each somata object, and then the mean of all somata object-wise scores in the image were calculated to create a FOV-wise score.

**Supplementary Figure 6. Example images showing a demarcated cell death event in image analysis.** Cell death was assumed where there was >10% drop of mean fluorescence in successive images, as seen at day 33 in this culture (human laminin, NB media and 1×10^5^ seeding density). Scale bar = 200𝜇m.

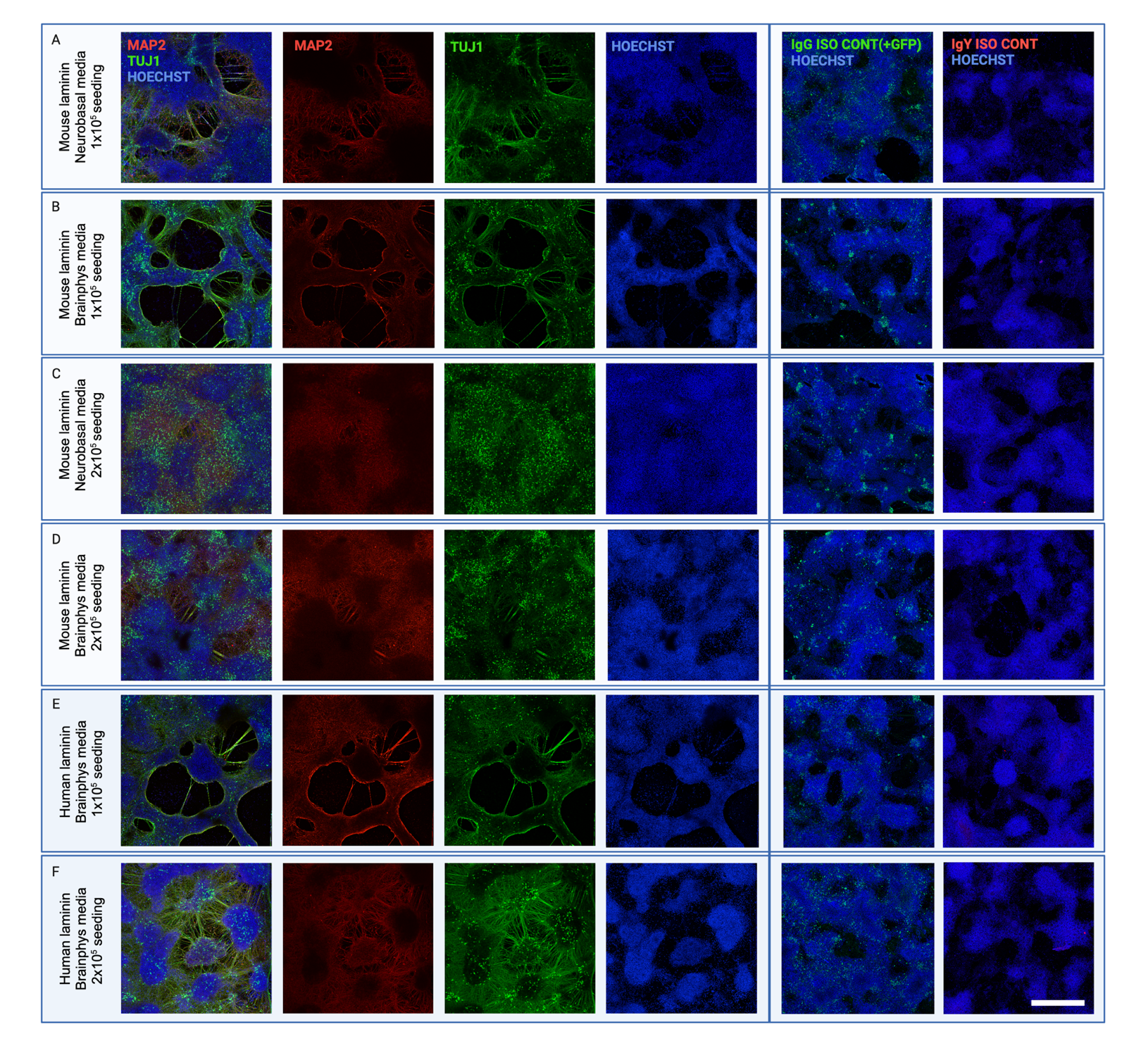

**Supplementary Figure 7. Morphological and molecular characterisation of cells post-experimentation at Day 40.** (**A-D)** Immunostained cultures displayed different levels of neurite blebbing and fragmentation across conditions after longitudinal imaging. Cells plated on mouse laminin with Neurobasal^TM^ (NB) or Brainphys^TM^ Imaging (BPI) media displayed extensive detachment and disorganisation. (**E-F)** Cells plated on human laminin with BPI media displayed moderate neurite fragmentation while still retaining major cytoskeletal structures. This is consistent with the extended lifespan observed in BPI media and human laminin-plated cultures, which underwent the last death event of all conditions (see Figure 3a). All cultures stained positive for neuronal markers class III beta tubulin (TUJ1; green) and microtubule-associated protein 2 (MAP2; red). Negative controls included rabbit IgG isotype control matched to TUJ1 primary (IgG Iso Cont; green; note that visible green fluorescence corresponds to residual GFP tagging on neurons) and chicken IgY isotype control matched to MAP2 primary (IgY Iso Cont; red). Cells were not able to be imaged in NB media x human laminin conditions due to complete detachment. Of note, immunostained images were acquired after defined death events and thus do not represent morphological and network features present throughout culture lifespan. Scale bar = 600𝜇m.

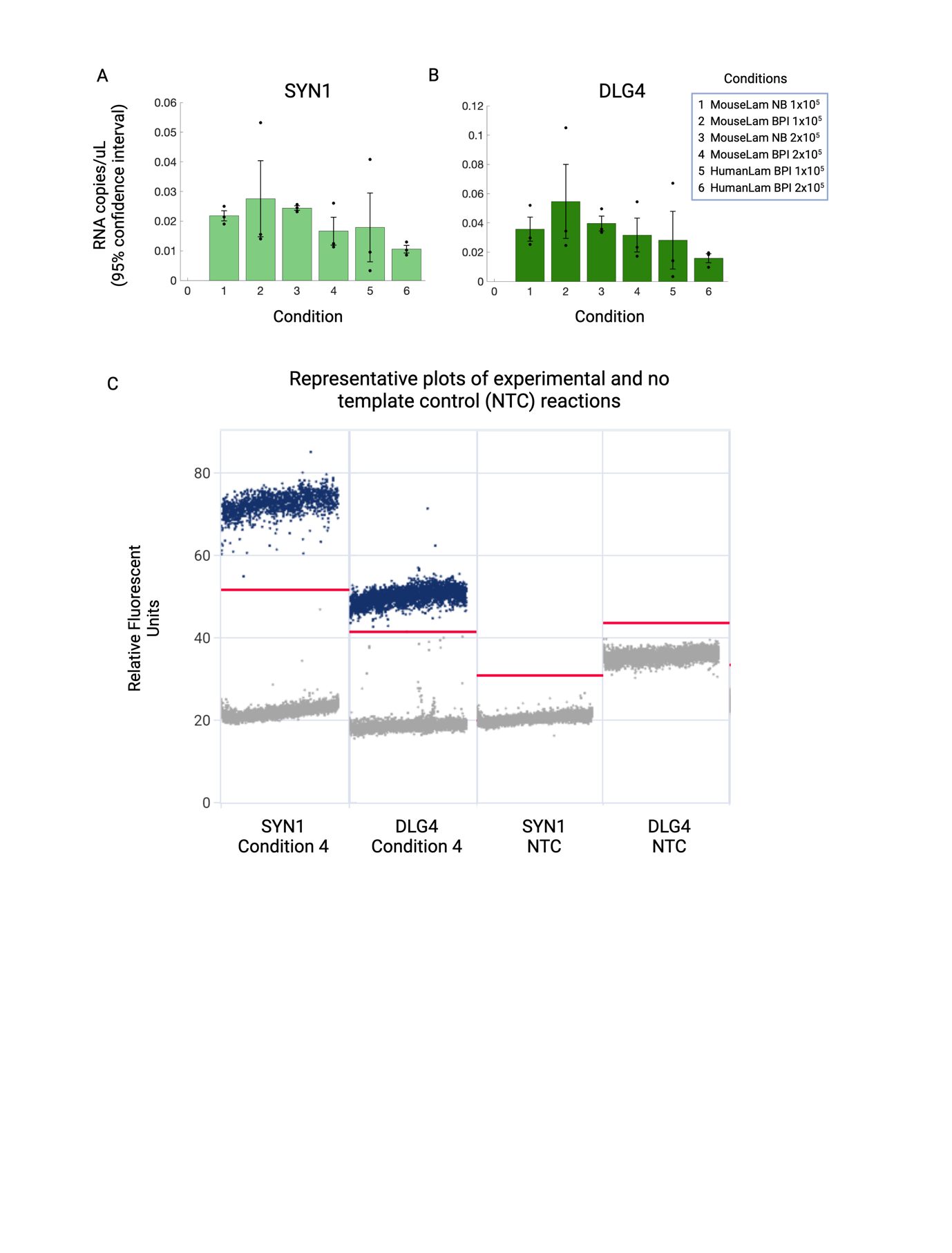

**Supplementary Figure 8. Digital polymerase chain reaction (dPCR) absolute quantification of SYN1 and DLG4 levels across culturing conditions at Day 42**. **(A,B)** *SYN1* and *DLG4* expression was present across all analysed conditions. Error bars denote standard error of the mean. Student’s t-test revealed no significant differences between conditions. Gene quantification was not conducted for NB media x human laminin conditions due to insufficient RNA concentration resulting from early death events. **(C)** Example plots of gene absolute quantification for condition 4, showing clear distinction between positive (blue) and negative (grey) partitions, and no template controls (NTCs), showing all negative partitions. Red lines denote thresholds automatically set by QIAcuity software. Data generated from 3 experimental plates (n = 3 per condition).

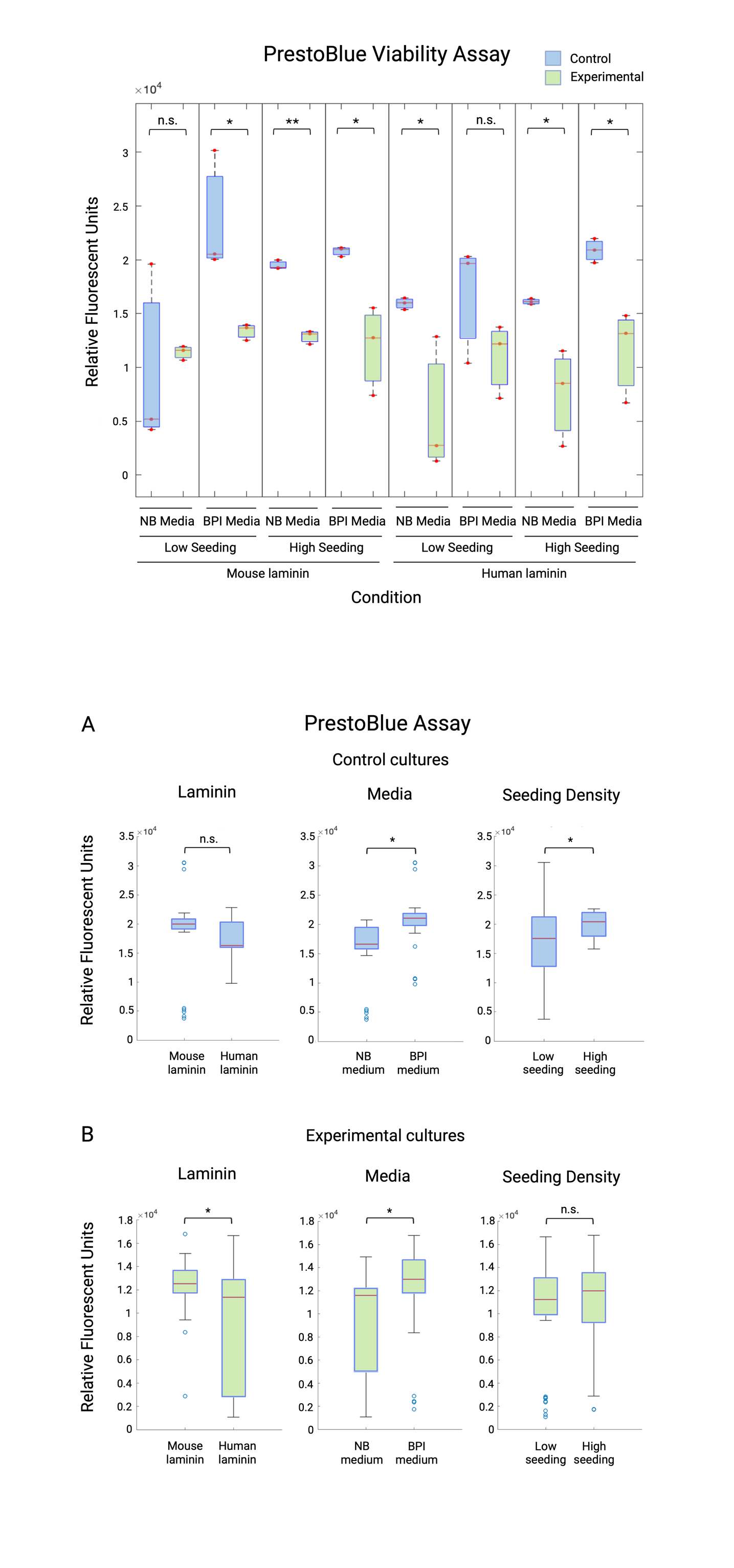

**Supplementary Figure 9. PrestoBlue assay reveals effects of laminin type, media, and cell seeding density on viability. (A)** Regression models with a within-subject factor for each condition compared the effect of laminin type, culture media and seeding density on viability in control cultures. Viability was higher in Brainphys^TM^ Imaging (BPI) media conditions relative to Neurobasal^TM^ (NB), and high seeding density (2×10^5^ cells per cm^2^), relative to low (1×10^5^ cells per cm^2^), but no difference was observed between laminin types. (**B)** Viability in experimental cultures was higher in mouse laminin conditions relative to human laminin, and BPI media relative to NB media, but no difference was observed between seeding densities. Boxplots represent median and interquartile range, whiskers denote upper and lower values, blue points represent outliers that fall more than 1.5 times the interquartile range outside boxes. Individual data points represent averaged triplicates from the same assay well. Data generated from 3 experimental plates (n = 3 per condition) and 3 control plates (n = 3 per condition). *p ≤ 0.05; **p < 0.0001; n.s. = no significant difference.

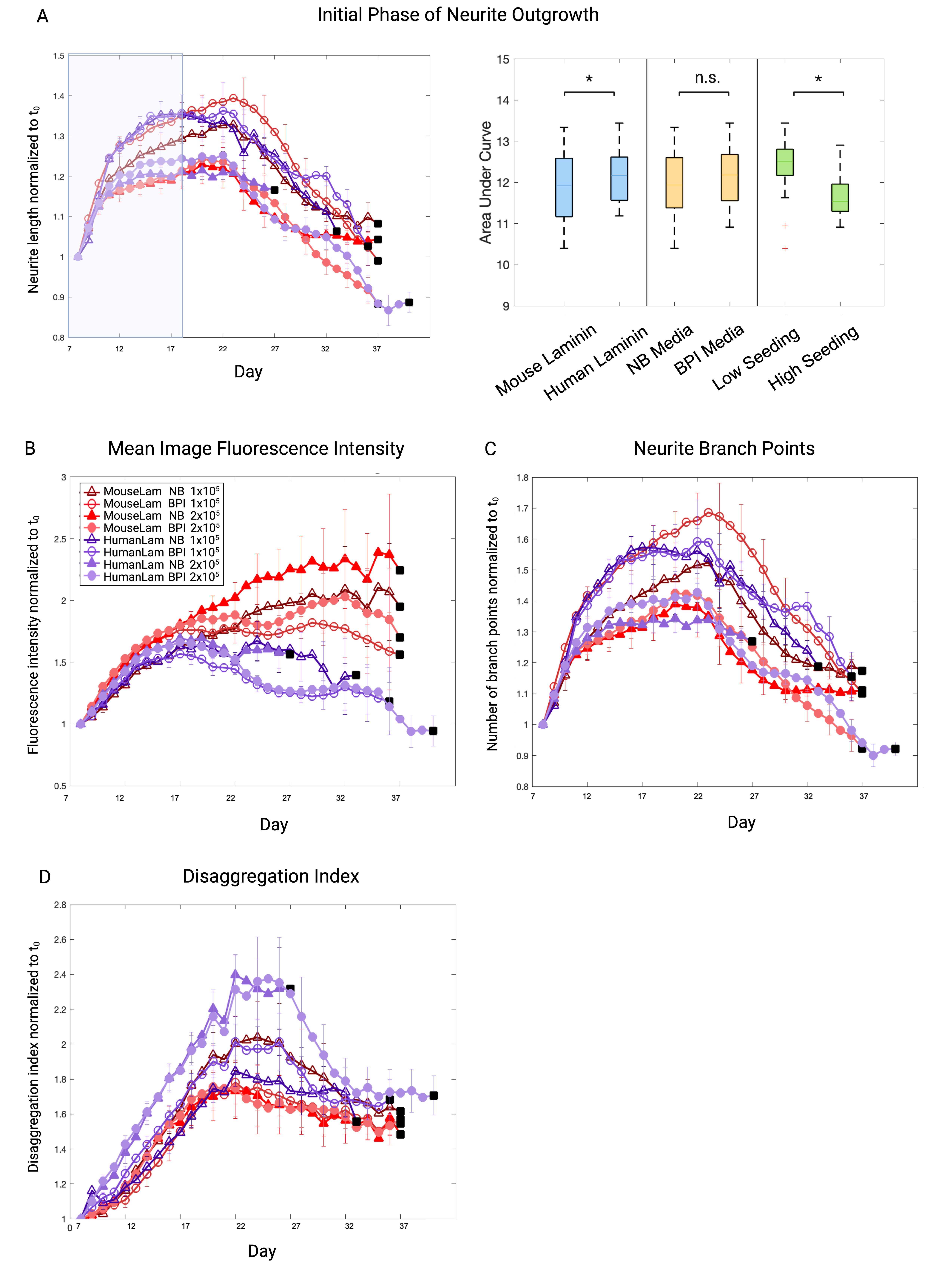
 **Supplementary Figure 10. Timewise plots of morphological metrics across 8 conditions, normalised to initial timepoint (t_0_). (A)** Analysis of neurite length in the initial growth period (Day 7-11) revealed higher neurite outgrowth in human laminin relative to mouse laminin conditions, and high seeding density (2×10^5^ cells per cm^2^) relative to lower density (1×10^5^ cells per cm^2^). No significant difference was observed between media. **(B)** Mean fluorescence intensity displayed an upward trend in mouse laminin and BPI conditions, likely due to bright somata clustering and thicker neurite structures. (**C)** The number of branch points was highest in BPI and mouse laminin conditions, indicating higher neurite complexity. (**D)** The Disaggregation Index showed more uniform somata patterns in BPI medium conditions. Plot datapoints represent mean of wells, and error bars represent standard error of the mean. Black squares demarcate cell death as defined by a >10% reduction in cell fluorescence between successive timepoints.

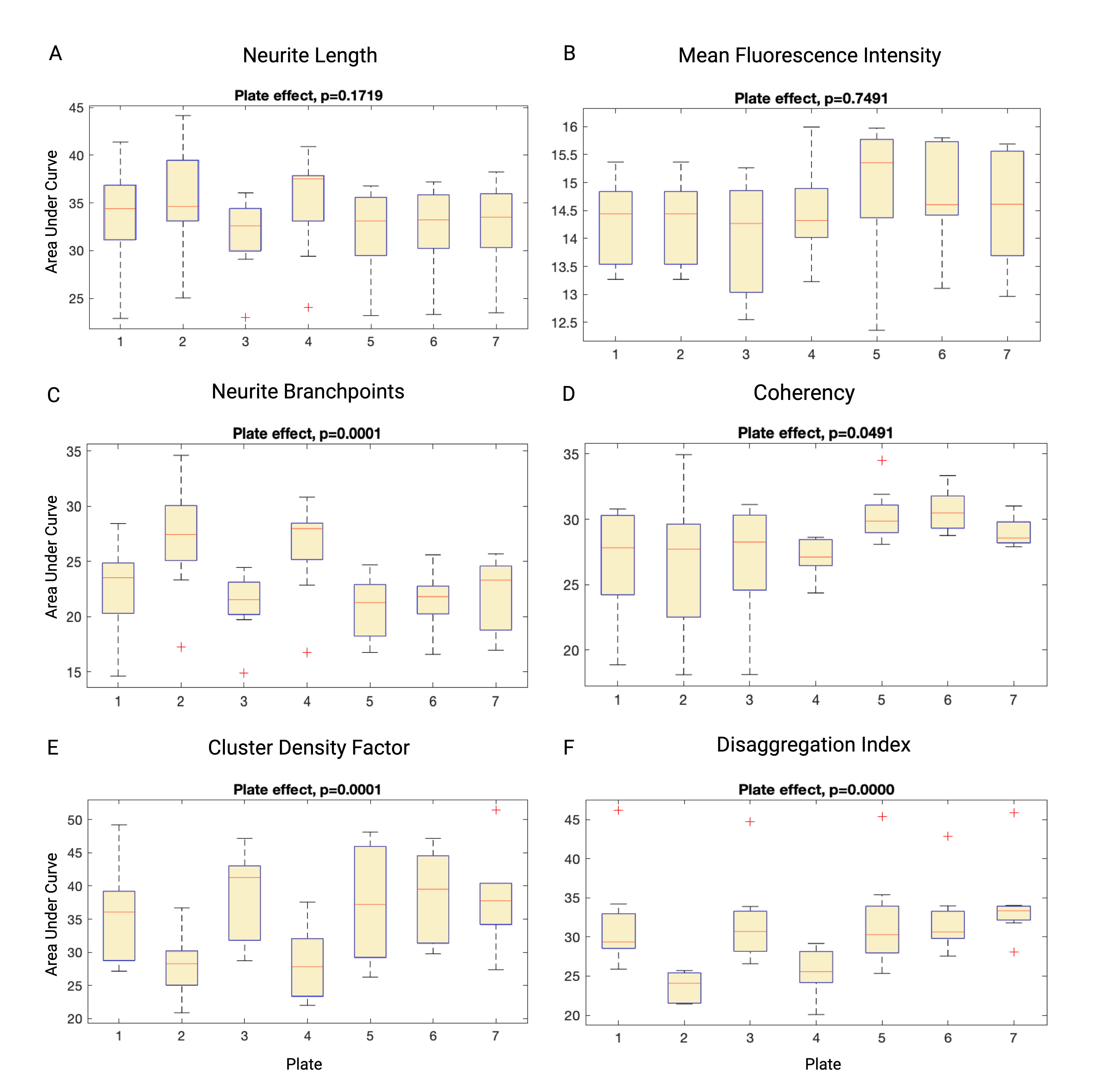

**Supplementary Figure 11. Comparison across plates for each morphological metric. (A-D)** No significant differences between plates were found for neurite length, mean fluorescence intensity, or neurite branch points. (**E-G)** Significant platewise differences were found for coherency, CDF and DI. Boxplots represent median and interquartile range, whiskers represent upper and lower values.

| **Metric** | **Laminin** | **Culture media** | **Seeding density** | **Laminin × media** | **Laminin × seeding density** | **Media × seeding density** |
| --- | --- | --- | --- | --- | --- | --- |
| **Fluorescence**  **Morphological and network metrics**  ***Neurite length***  ***Branch points***  ***DI***    ***CDF***  ***Coherency***  **PrestoBlue Viability Analysis**  ***Control cultures***  ***Experimental . cultures*** | t = -10.60, *p* < 0.0001*  t = -5.67, *p* < 0.0001**  t = -7.84, *p* < 0.0001**  t = 1.44, *p* = 0.2149  t = -1.55, *p* = 0.1908  t = -7.04, *p* < 0.0001**  t = -0.89, *p* = 0.3775  t = -3.84, *p* = 0.0003* | t = 1.66, *p* = 0.1708  t = 6.71, *p* < 0.0001**  t = 8.27, *p* < 0.0001**  t = 4.52, *p* < 0.0001**  t = 8.20, *p* < 0.0001**  t =6.77, *p* < 0.0001**  t = 4.83, *p* < 0.0001**  t = 2.89, *p* = 0.0052* | t = 1.15, *p* = 0.3389  t = -4.44, *p* < 0.0001**  t = -1.47, *p* = 0.2154  t = 1.98, *p* = 0.1144  t = 3.21, *p* = 0.0062*  t = 2.78, *p* = 0.0190*  t = 2.62, *p* = 0.0108*  t = 0.69, *p* = 0.4929 | t = 5.15, *p* < 0.0001**  t = 5.10, *p* < 0.0001**  t = 0.73, *p* ­­­= 0.4718  t = -0.30, *p* = 0.8353  t = -0.82, *p* = 0.5376  t = 1.88, p = 0.0666  -  - | t = 1.85, *p* = 0.1217  t = 1.86, *p* = 0.8025  t = 0.50, *p* = 0.6205  t = -0.05, *p*= 0.9631  t = 0.54, *p* = 0.7376  t = -0.99, p = 0.3271  -  - | t = -2.05, *p* = 0.1037  t = -0.46, *p* = 0.7745  t = 0.28, *p* = 0.7783  t = 1.96, *p*= 0.1069  t = -0.29, *p* = 0.8183  t = -0.12, p = 0.9067  -  - |

**Supplementary Table 1. Regression model results for morphological and viability data, adjusted with Benjamini–Hochberg correction.** Fluorescence and morphological metrics derived from area under curve of timewise plots, and PrestoBlue viability analysis derived from normalised relative fluorescent units. *p ≤ 0.05; **p < 0.0001. CDF = Cluster Density Factor, DI = Disaggregation Index.

| **Condition** | **Cluster Density Factor** | | **Neurite Coherency** | | **Summed rank** | **Overall network score** | **Summed mean AUC** |
| --- | --- | --- | --- | --- | --- | --- | --- |
|  | Mean AUC | Rank | Mean AUC | Rank |  |  |  |
| Mouse laminin  NB media  Low seeding | 29.20 | 6 | 26.12 | 5 | 11 | 3 | 55.32 |
| Mouse laminin  BPI media  Low seeding | 35.57 | 3 | 26.50 | 4 | 7 | 2 | 62.07 |
| Mouse laminin  NB media  High seeding | 34.06 | 4 | 28.00 | 3 | 7 | 2 | 62.06 |
| Mouse laminin  BPI media  High seeding | 36.37 | 2 | 28.45 | 1 | 3 | 1 | 64.82 |
| Human Laminin  NB media  Low seeding | 20.86 | 8 | 21.67 | 7 | 15 | 4 | 42.53 |
| Human Laminin  BPI media  Low seeding | 31.89 | 5 | 24.57 | 6 | 11 | 3 | 56.46 |
| Human Laminin  NB media  High seeding | 24.45 | 7 | 17.38 | 8 | 15 | 4 | 41.83 |
| Human Laminin  BPI media  High seeding | 43.14 | 1 | 28.43 | 2 | 3 | 1 | 71.57 |

**Supplementary Table 2. Calculation of overall network score based on Cluster Density Factor and neurite coherency.** Mean AUC was ranked across all conditions for each of the two metrics, and their rank was added to form a summed rank. Summed ranks were then inversely ranked to determine overall network scores; with lower summed ranks denoting higher network scores and vice versa. The summed mean AUC across both metrics was also calculated.
